# Knowledge Innovation Ecosystem for the Promotion of User-Centre Health Innovations: Living Lab Methodology and Lessons Learned Through the Proposal of Standard Good Practices

**DOI:** 10.1101/2024.01.17.573578

**Authors:** Natacha Rosa, Sofia Leite, Juliana Alves, Angela Carvalho, Diana Oliveira, Flavia Santos, Barbara Macedo, Hugo Prazeres

**Affiliations:** i3S - Instituto de Investigação e Inovação em Saúde, Universidade do Porto, Rua Alfredo Allen, 208, 4200-135 Porto, Portugal; INEB - Instituto de Engenharia Biomédica, Universidade do Porto, Rua Alfredo Allen, 208, 4200-135 Porto, Portugal; IBMC - Instituto de Investigação e Inovação em Saúde, Universidade do Porto, Rua Alfredo Allen, 208, 4200-135 Porto, Portugal; IPATIMUP – Instituto de Patologia e Imunologia Molecular da Universidade do Porto, Rua Alfredo Allen, 208, 4200-135 Porto, Portugal

**Keywords:** Living Labs, Healthcare, Good practices, innovation

## Abstract

Living Labs, experiencing a global surge in popularity over the past years, demands standardized guidance through the development of widely accepted good practices. While challenging due to the complex and evolving nature of Living Labs, this task remains essential. These knowledge innovation ecosystems facilitate a diverse array of interconnected and interacting end-users and stakeholder partners who engage collaboratively to co-create, embed, and/or leverage end-user-centric breakthroughs at one or more innovation phases within a real-world context. Based on the development of six Living Labs in the health domain, this study proposes a more general yet critical set of Living Labs’ good practices, emphasizing the importance of strong initial marketing and promotion strategies for Living Labs’ open calls, enforcing gender equality, carefully selecting stakeholders, devising and implementing effective framework strategies for end-user engagement and value creation, ensuring value creation for all Living Labs partners, prolonging the long-term viability of the Living Lab project, promoting and disseminating impactful actions and results, fostering environmental sustainability, and processing results data for Living Lab performance evaluation.

## 1. Introduction

The global economy is moving towards a knowledge-intensive model and innovation was identified as a key driver for economic and social growth [1]. However, the ability to develop impactful breakthrough services and products has far-reaching implications [2] and it is not a straightforward process [3]. This is due to the growing complexity and rapidly changing market environment [4] which consequently leads to a demanding and hampered concept associated with innovation success, such as technical uniqueness, competitive advantage, diversity of market offering, protection of a market position, and profitability [5]. As a consequence, there is around 40% rate (in contrast to the popular invalidated belief of being around 80%) in innovative product failure, as the percentage of new products introduced to the market that then fail to meet the commercial objectives of the business unit that launched the product [5]. The rates of idea failure are impractical to quantify due to the unavailable concrete factual data, however, they are expected to be significantly higher than the product failure rate. This scenario urges innovators to a constant flow of ideas while competing through emergent technologies and fast new product development [6].

The term Living Labs (LLs) was first proposed in the United States in the 1990s by Lasher *et al.* [7]. However, it wasn’t until 2006 that this methodology gained a significant increase in Europe by the European Commission [8]. This KIE was introduced in an attempt to overcome the gap between knowledge production and innovation commercial success. This so-called “European Paradox”, through policy measures, advanced, coordinated and promoted a common European innovation system [1, 4]. The LLs have been applied around the globe, to generate impactful innovation within and suited to real-life problems and contexts [8]. These systems are associated with different conceptualizations and there is no general agreement in terms of their standard definition [9, 10]. However, based on our experience and understanding, we define LLs as a knowledge innovation ecosystem (KIE) of diverse interconnected and interacting end-users and other stakeholder partners which engage through keenness, trustful relations building and knowledge sharing to co-create, entrench and/or leverage end-user-centred breakthroughs at one or more innovation phases and in a real-life driven-context (Figure 1). Similarly to Yasuoka *et al.* [9] suggestion, in our definition, the LLs collaborators are regarded as "partners that create a service together" rather than mere subjects or targets for experiments. Also, the involvement of end-users in the innovation process and the evaluation of the solutions in a real-world setting, are considered the two core concepts that make LLs distinguishable [11, 12] and which will determine the innovation impact and consequently on whether and how society adopts them [13]. As stated by Logghe and Schuurman [4] end-users ideas, experiences, and knowledge, as well as their daily needs, are essential inputs for the starting point in innovation, as well as, to enhancement and ‘shaping’ of products and services to satisfy and fit the specific market demands whereas consumers have a higher willingness to pay for a product or service that perfectly satisfies their personal needs. Also, the LL’s ability to allow the solutions’ testing with target end-users at key innovation phases is expected to boost the societal impact of the achieved technical breakthrough, in terms of product acceptance, adoption and delivered value [11, 14]. Consequently, the resource to the LLs methodology will reduce market risk in the launch of innovative offerings while improving the return on investment and time to market [6].

**Figure 1.**
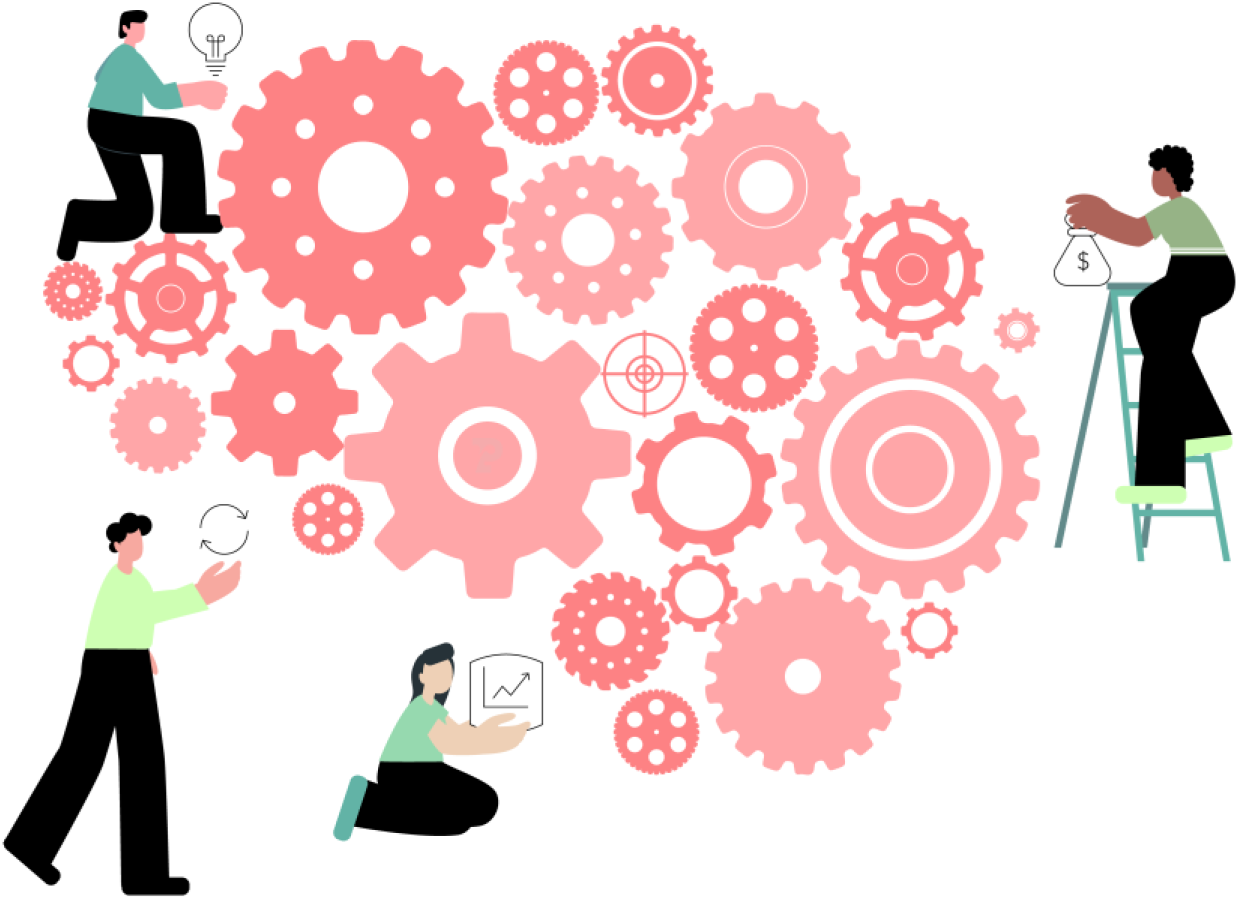
Schematic representation of the LL definition. The knowledge innovation ecosystem, represented by the brain-shaped gear wheels, enables end-user-centred innovation co-creation, entrenching and/or leveraging through the interconnection and collaboration of the LL’s different role participants

According to recent data from the pan-European network ENoLL (community aiming to promote and enhance the LLs concept globally) [15], there are more than 480 historically recognized LLs over 14 years. In Portugal, in recent years, there was a registered increase in the adhesion to and experimentation with LLs for the leveraging of innovations in many sectors and diverse fields of application from open schooling methodology (SALL schools as Living Labs [16]) to urban electric mobility (RENER Living Lab). However, most cases ended-up being sporadic actions with no indication of continuity. From the fourteen Portuguese LLs accredited by EnoLL in 2013, as stated in the study developed by Álvaro de Oliveira and David Amaral de Brito [17], currently only the Smart Rural Living Lab is indicated as being active.

Despite the potential demonstrated by the LL approach, these KIEs are challenging schemes characterized by multi-actor, multi-function, and multi-scalar dynamics [18]. Also, the guidelines methodology, framework or practices are neither backed-up by consistent scientific research nor by supporting theories on good practice of innovation and technology transfer and interaction between the different actors in this context [1, 18, 19]. Hence, in this article, the Resolve-Health program, based at the Institute of Health Innovation and Research (i3s in Portuguese *Instituto de Investigação e Inovação em Saúde*) LL initiative will be considered as an exploratory case study. In particular, the six LLs’ initiatives in which all the authors participated from 2022 to 2023, are the study focus of this article. For the interest of clarity the considered LLs are in the health-related field, namely, Patient-Centered Cancer Living Lab Pediatric Oncology; The Brain Living Lab; Remote Monitoring Living Lab; Digital Health Validation Living Lab; The Microbiome in Agro-Food and Health Biotech Living Lab and Innovations in Animal Health Living Lab. In this article, common patterns and points of divergence in the considered LLs’ will be considered for the evaluation of the approaches adopted and the clarification and discussion of their impact on the KIE process. Therefore, the purpose of this article is to analyze from a technology transfer perspective the strategies adopted in these LLs’ and with that identify and propose a set of good practices for the successful implementation and evaluation of LLs. What we present in this article is an attempt to propose a set of good practices to be recognized or used as a reference by the scientific community and by other LLs in the health domain field. Accordingly, qualitative and quantitative efficiency evaluation metrics for both the performance of LL processes and their broader impacts were also suggested. In the long term, this study also aims to encourage a more general adoption of this KIE.

The remainder of this paper is structured as follows. First, in Section 2, the methodologies adopted in the LLs are described. Section 3 is dedicated to the presenting of the results related to LLs’ participant teams and their innovations, collaborating stakeholders, engagement strategies, value created for all the LL’s partners, as well as, the outcomes dissemination strategies. The results obtained are discussed in Section 4, through the proposal of a set of nine good practices where the study ascertainment limitations and the result’s future implications are also addressed. Section five presents the conclusions.

## 2. Methods

The LL’s engaging roles are assembled in five distinct groups (inspired by Leminen and Westerlund [6] LLs actors’ classification group), namely: (1) providers (that contribute with the core innovation, *i.e.* idea, technology and/or service and that can be at different evolution stage); (2) end-users (potential direct clientele of the provided innovation); (3) experts (professionals or *connaisseurs* that promote and leverage innovation in its evolution stage through power, technical specialized know-how, process and relationship but are not their potential direct clientele); (4) hosts (organizations or ecosystems that provide support for the development of the LLs in terms of virtual or physical space with or without technical support material, *e.g.* projectors, tables and chairs, coffee machine for coffee breaks etc., and that could also provide access to stakeholders) and (5) liaisons (officers which interconnect and articulate the KIE different intervenient as well as coordinate and define/co-define the activities of mutual concern). The end-users and the experts are both stakeholders. However, given the specificity and relevance associated with the end-user role in the LLs system, a category to differentiate non-end-user stakeholders and end-user stakeholders was created.

### 2.1. LLs case studies and providers

Under the premise of obtaining a representative, deeper and reliable understanding of this complex modern KIE phenomenon and consequently developing an efficient analysis and evaluation of the LL impact [8, 20], a case research study approach, was considered. To compare and contrast its implementation approaches and outcomes, a multiple case research strategy on six LLs was adopted in this article: Patient-Centered Cancer Living Lab Pediatric Oncology (PCCLL); The Brain Living Lab (TheBrainLL); Remote Monitoring Living Lab (RemotMonitLab); Digital Health Validation Living Lab (DigitalHealthLL); The Microbiome in Agro-Food and Health Biotech Living Lab (MicrobiomeLab) and Innovations in Animal Health Living Lab (AnimalHealthLL). These LL initiatives aimed to provide leveraging and management tools to early-stage projects and spin-offs in Health Sciences to transform innovative knowledge into business ventures and value creation. Hence, the program was not extended to large players and SMEs as innovative solution providers as is often suggested in the literature [14]. These LLs were selected as case studies because they represent successful examples where different stakeholders collaboratively addressed specific innovation challenges and where end-users had an active and determinate contribution in the design, validation and testing of the innovative technologies and services. Also, the continued duration of the LLs allowed for the gathering of a considerable amount of information.

Resolve-Health action started by setting an international individual open calls for each of the LLs with information regarding their scope and type of innovations supported (Table 1), as well as, the technical support provided namely completion of proof-of-concept (PoC), validation of prototypes and product development with feedback from end-users and within a living lab context. Besides the technical support provided by the LLs’ the call also announced financial support valued between 5€K to 20€K and adapted to each specific project’s needs. The open calls were disseminated through the Resolve-Health webpage and social media (such as LinkedIn).

**Table 1.** LLs open calls information in terms of the acronym used, innovation supported by the LL, call promotional time and the correspondent collaborative host.

| LLs |  | Host | Promotional time (weeks) |
| --- | --- | --- | --- |
| Abbreviation | Innovation supported |  |  |
| PCCLL | Solutions (preference given to e-health) to improve the health and quality of life of pediatric oncology patients and their families during patient journeys | Associação Acreditar [21] | 6 |
| TheBrainLL | Pharmacological therapies for the diagnosis, monitoring, treatment and rehabilitation of neuropsychiatric, neurological, neurodegenerative and neurodevelopmental disorders | Centro CEREBRO [22] | 5 |
| RemotMonitLab | Technology-based solutions for mental health tailored support | Neurobios [23] | 2 |
| DigitalHealthLL | Digital tool solutions for diagnosis, monitoring, treatment and rehabilitation in HealthCare | CUFF [24] | 2 |
| MicrobiomeLab | Microbiome-based solutions to address new and existing threats to food security, nutrition, health and agri-food systems' sustainability | Biocant [25] | 3 |
| AnimalHealthLL | Small domestic animals' health-based solutions for the promotion of vet medicine new technologies, therapies and treatments | Onevet - Hospital Veterinário do Porto [26] | 2 |

The LLs’ applied innovations information was collected from an online application form filling (form data can be seen in the Supplementary Material, Supplementary Figure 1). The submission of additional documentation was required to be sent by e-mail to the Resolve-Health team. This procedure was adopted to define and guarantee a standardized and automatic delivery of information regarding the applied innovations’ content. The innovations were classified according to the Technology Readiness Level (TRL) which is a systematic metric/measurement system that supports assessments of the maturity of a particular technology and the consistent comparison of maturity between different types of technology [27]. The evaluation process involved two phases, the first was based on the information provided in the application form and the second was based on an interview. In the first evaluation phase, the innovations that did not fill the minimum LLs’ requirements (*e.g.* out of the scope with the LL it applied for and all the other LLs available, initiatives which were in the early ideation process or with undefined goals) did not proceed in the selection process. The innovations selected for the second phase were subjected to an interview by the Resolve-Health team and in collaboration with the corresponding LL host manager. After the providers’ short presentation through a pitch, the interviewers adjusted their questions with flexibility on-site based on the information provided, their answers and interactions with interviewees. These interviews allowed the clarification of details regarding the technologies/services and also to manage the applicant’s expectations towards their participation in the LL. The interviews were conducted virtually using the video-based platform ZOOM Cloud Meetings version 5.14.0 (Zoom Video Communications, Inc., San Jose, California) with conversations typically lasting up to 1 hour. The innovations applied were afterwards evaluated by a multidisciplinary jury, including the RESOLVE Team and experts, gathering academic, clinical and business perspectives and the corresponding collaborative LL host manager and/or team. The selection of the participant’s innovative projects was based on the scope of LL action and according to the following criteria: potential and social impact (20%), feasibility in the context of each living lab (20%), uniqueness of the idea/concept (20%), team experience and connection to the project (20%) and market potential and scalability (20%). The LL’s screening parameters customization strategy was adopted to guarantee an appropriate and efficient innovation project selection so the experience could be enriched/fostered for both the participant and host ecosystem.

### 2.2. Stakeholders

Stakeholders were carefully considered and invited to participate in the LLs network. Their selection was based on the innovation requirements and the stakeholder’s availability, interest and level of contribution. Involved stakeholrs references were obtained from successful previous engagement with the Resolve-Health activities and defrom the hosts which provided access to internal and external link stakeholders. The following terminology was considered to classify the participant stakeholders (end-user or expert) in this study:

- ‘Government’ is a stakeholder from any foreign central, regional or local government department, agency, or other entity performing governmental functions which includes governmental corporations or their separate business units (definition adapted from [28]). This term does not comprise stakeholders from research institutions or organizations, educational organizations or medical organizational entities that are performing governmental functions.
- ‘Medical’ is a stakeholder from public and private health organizations (*e.g.* health centres and hospitals);
- ‘School’ is a stakeholder from any educational organization (private or public), level (*e.g.* secondary school, tertiary higher education) and position (*e.g.* students, professors and school technicians);
- ‘Research’ as stakeholders from R&D institutes or centres;
- ‘NGO’ (non-governmental organization which can also be denoted as "non-commercial organization", "non-profit organization”, "non-profit association" or "organization without lucrative purpose") is a stakeholder from an organization that is established by the free will of the citizens who are associated on common career interests and/or other interests aiming to achieve shared civil, economic, social and cultural rights and not obtaining profits [29];
- ‘Company’ is a stakeholder from an entity that represents an individual or group of individuals who conduct commercial business practices to earn a profit;
- ‘Citizen’ is a stakeholder (or end-user) that is the public (or final) receiver of a product or service.

### 2.3. LLs methodology/approach

The LL’s activities were designed and selected based on the action research methodology. In this iterative approach, there is a circle of planning, action, and fact-finding about the result of the action [4, 30]. The proposed innovations development stage, the technical support still required, the end-users’ unmet needs, their variable engagement level as well as the feedback from all the participants (providers and stakeholders) were considered as the result and action strategy in the design and creation of each LL activity and session.

Considering that the LLs’ activities require confidential information to be shared, all the participants in the activities (liaisons, host collaborators, providers, and stakeholders) and the steering committee were obliged to sign a non-disclosure agreement (NDA). A consent to photograph and video capture form was also signed by all the participants.

### 2.4. LL partners’ feedback acquisition and assessment

The providers’ feedback on the LL experience and the sessions they participated in was obtained by answering a facultative online questionnaire with close and open questions. This questionnaire was given to each provider’s team member who participated in the LLs activities (*e.g.* PhD students, researchers, scientific coordinators, head of innovation, start-up CEO, etc.). The questionnaires were delivered online through the Microsoft Teams survey tool. The respondent’s motivation and answers quality were guaranteed by revealing to them that the data collection would be considered not only for the RESOLVE-HEALTHfinal project metrics but also for the development of a research study and consequently for publishing the results as a scientific article format. The surveys were all confidential to guarantee the respondent’s privacy and to make them comfortable in expressing their true opinion. The survey was composed of three sections. The first was a set of 15 closed questions based on a 7-point Likert scale [31] and were grouped into four distinguishable categories: relevance of mentorship and sessions, specification of actual needs, final gains, and general satisfaction (assessing the perception of quality, relevance, and satisfaction). The 7-point response spectrum offers a range of choices that allow respondents to pick the ‘exact’ one prefer rather than to pick some ‘nearby’ or ‘close’ option [32]. The scale ranged from 1=totally disagree to 7=fully agree. The second section asked the participants to identify the three most relevant sessions (including the specialization phase training sessions) that the program contemplated, per order of relevance and according to the characteristics and needs of their project. The third and final section, through an open format, encouraged the sharing of any additional remarks that they might find relevant about the LL program or any session in particular *(e.g.* strengths, weakness, etc.). Data for this study were collected from three of the LLs at the end of July 2023: PCCLL, TheBrainLL and RemotMonitLab. The survey was developed by a credited psychologist. The survey form data is available in Supplementary Figure 2 in the Supplementary Material.

A qualitative and quantitative evaluation of the KIE experiences from the facilitator’s perspective was obtained from both a questionnaire and one-on-one interviews. The liaisons were asked to report through a questionnaire specific quantitative and qualitative data (*e.g.* characteristics and number of projects that participated in the LL they were coordinating) and what they registered during field observation, informal conversations with stakeholders and enablers and also their opinion regarding the developed LL strengths, weakness and future improvements.

Feedback (namely contentment, frustrations and suggestions) from the stakeholders and hosts was noted down and perceived through casual conversations during the LL sessions and activities.

Highly structured quantitative and qualitative key performance indicators were defined to evaluate the providers, stakeholders, value creation and results dissemination.

### 2.5. LinkedIn data

The number of followers represents the cumulative number of followers to the Resolve-Health [33]. The Resolve-Health engagement rate is a calculation of the number of likes, clicks, comments and shares on the Resolve-Health updates, divided by the impressions in the last seven days [33]. Page views correspond to the total number of page views for the different sections of the Resolve-Health page over a maximum period of 12 months [33]. The unique page visitors are the estimated number of unduplicated individuals who visited the LinkedIn Resolve-Health website [34] over twelve months [33, 35]. In this study, it was assumed that the individuals who visited the LinkedIn website were a result of the post published. Since this was considered as the sum of unique page visitors in the consecutive periods between the publishing of a new post with information regarding the LL activities and until another new post was published with other information about the Resolve-Health that number was not related to the LL.

### 2.6. Statistical analysis

All the Statistical analyses were performed with Microsoft Excel Software (Microsoft Corporation, 2018). The average values were calculated by adding all the data values and dividing this sum by the number of values. The deviations between the data values and the mean correspond to the spread or standard deviation in the data [36]. The average and standard deviation values were obtained from the excel functions *=AVERAGE(Range)* and *=STDEV.P(Range)*, respectively, where *Range* corresponds to the range of values.

## 3. Results

### 3.1. Providers

The percentage distribution of the participant innovations per field of application is presented in Figure 2.

**Figure 2.**
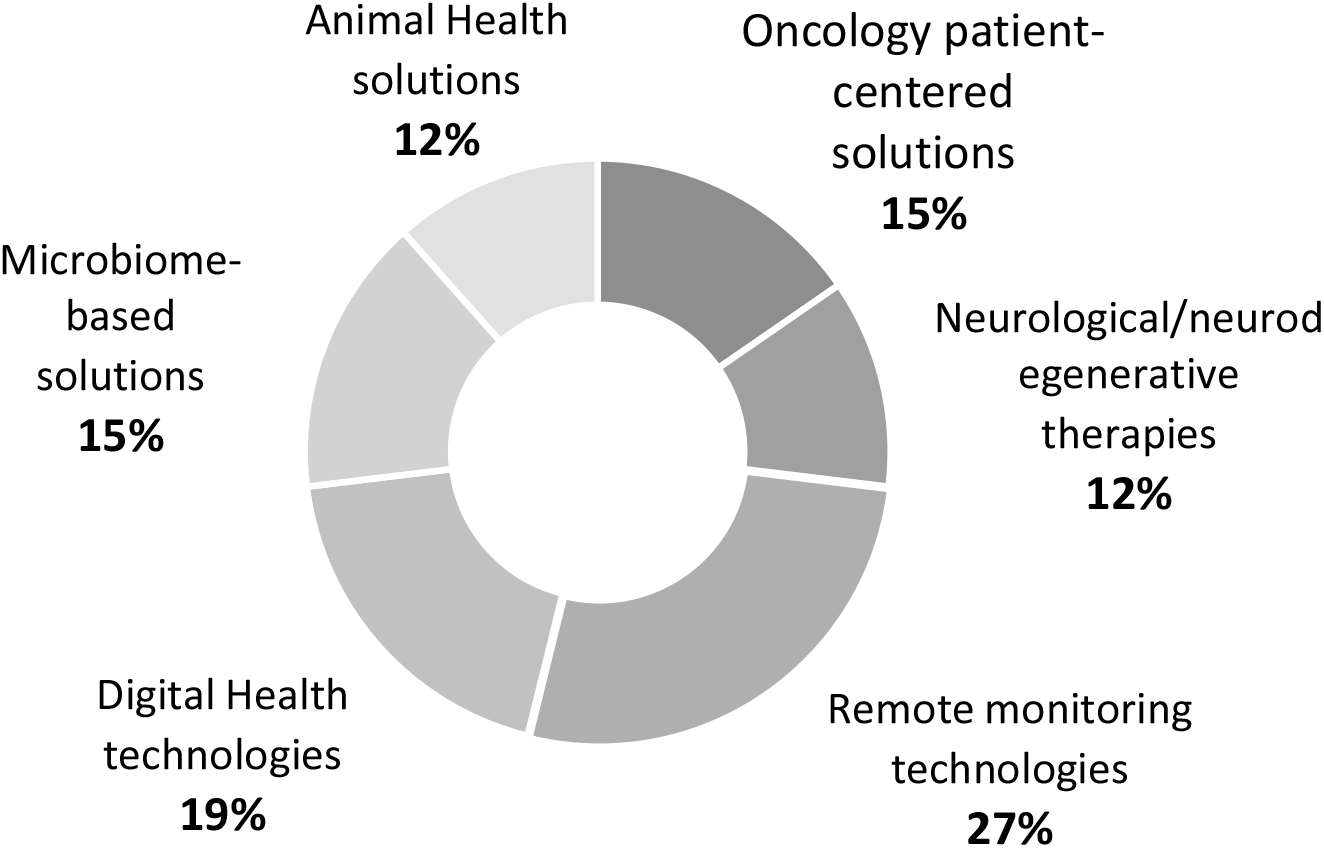
Percentage distribution of the participant project in the total LL per innovation filed of application

It is possible to verify, from Figure 2, that there is a higher number of innovation projects in the technology IT field with 27% and 19% in the remote monitoring and digital health area, respectively. Both microbiome-based and oncology patient-centred solutions correspond to 15% of all total innovation projects participating in the LLs. The projects in the animal health and neurological/neurodegenerative therapy field showed a lower number of participant projects. For more information, a detailed description of the participant projects, their TRL and technical support obtained per LL are presented in Table 2.

**Table 2.**
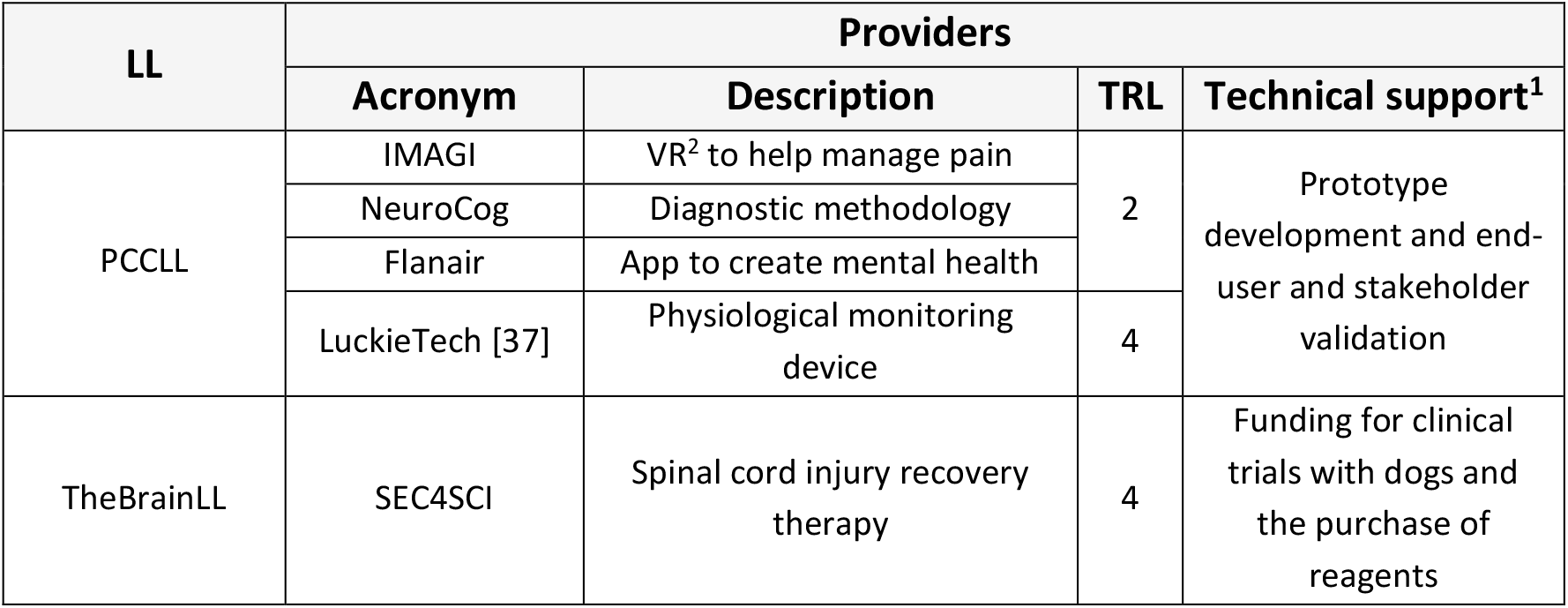
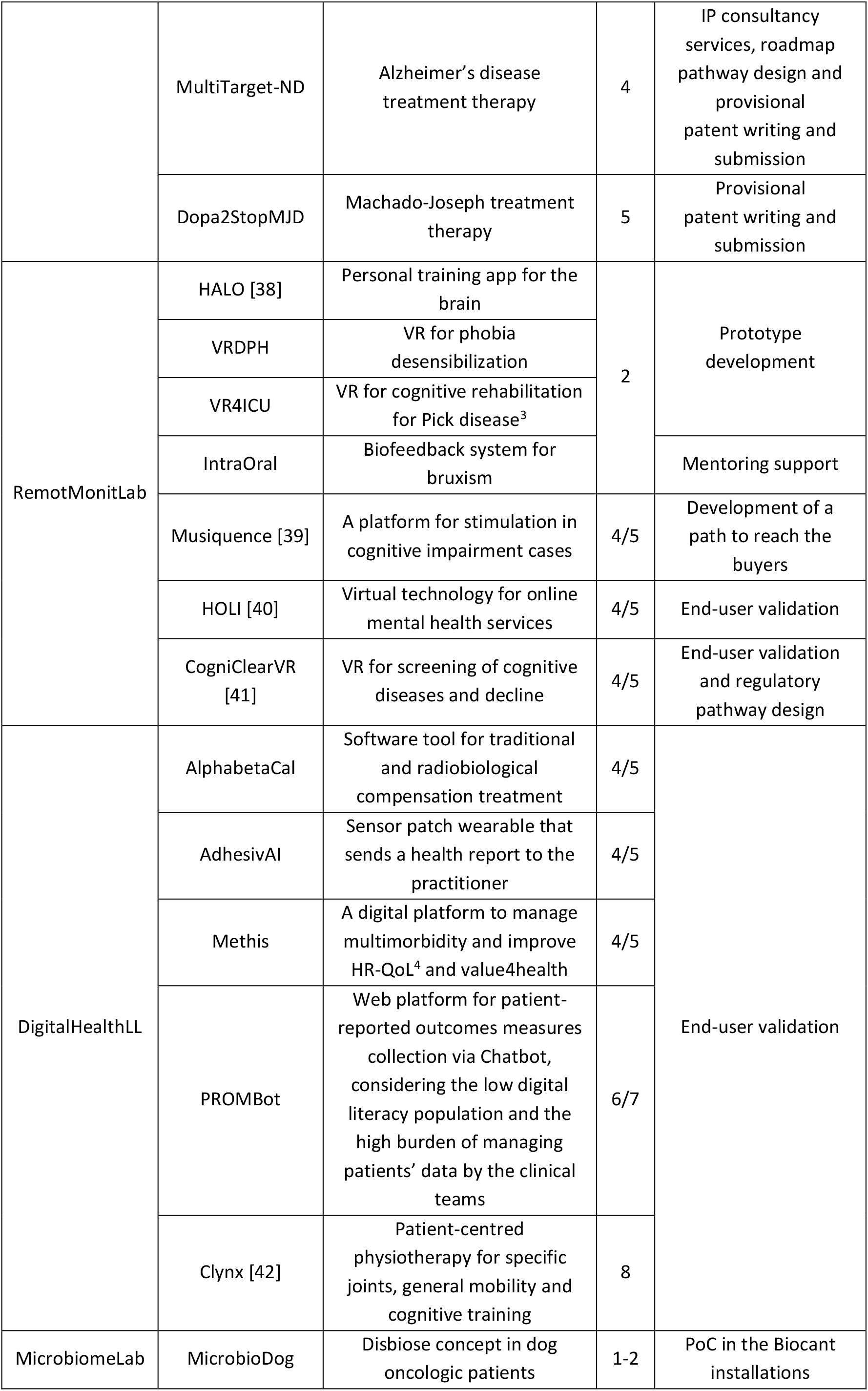
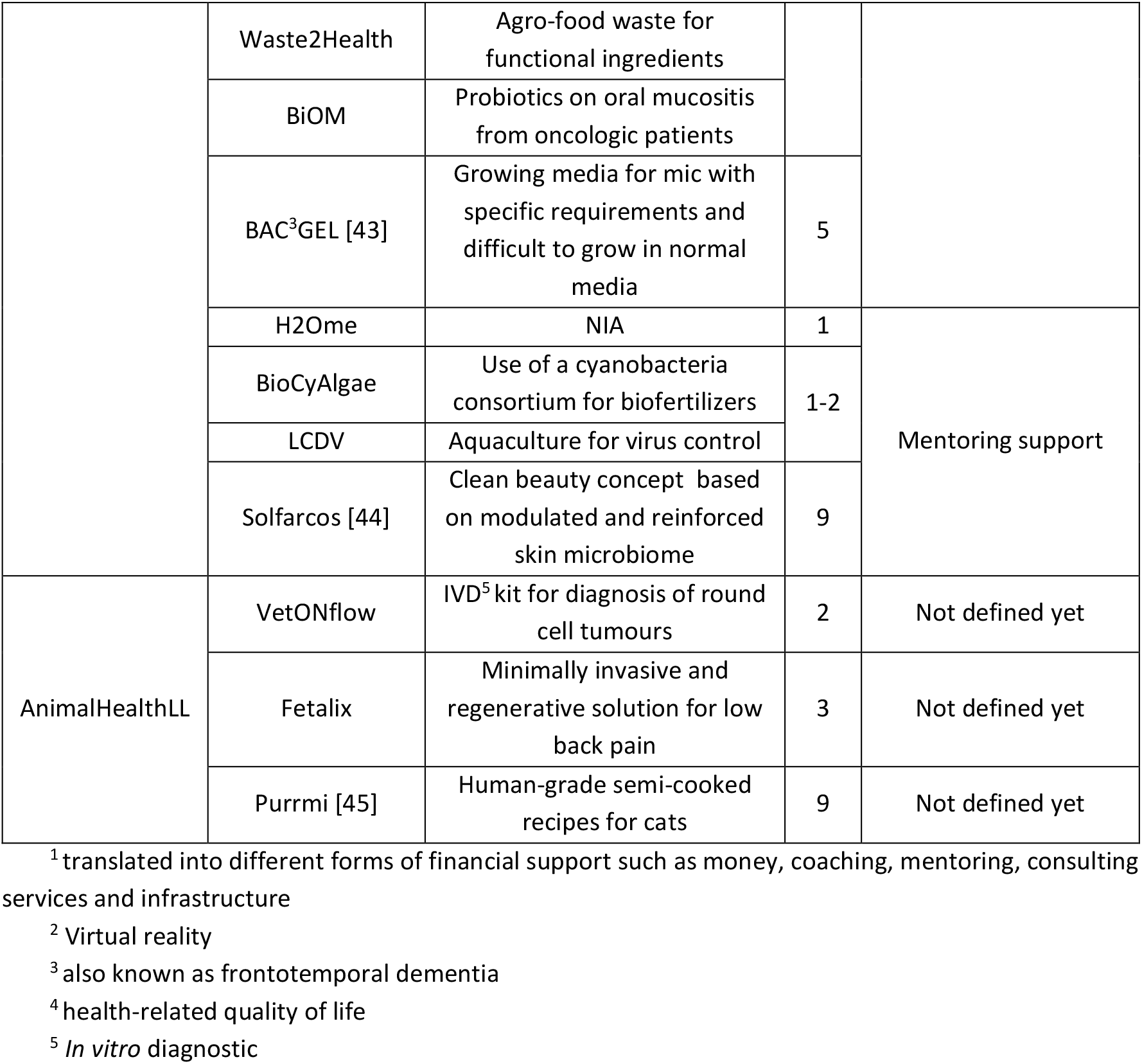
Correlation between the LLs’ and the selected innovation projects with information about their acronym, short description, technology readiness levels (TRL) and technical support obtained.

From Table 2, it is possible to verify that the KIE-supported innovations varied in terms of the type of innovation; TRL and technical support obtained. PCCL supported innovations from both the computer and medical technology fields. TheBrainLL supported only pharmaceutical solutions with a 4/5 TRL. RemotMonitLab and DigitalHealthLL mainly supported computer technology solutions. MicrobiomeLab supported biotechnology solutions and which TRL differ from 1 to 9. The AnimalHealthLL supported medical technology, pharmaceutical and animal nutrition solutions with TRL from 2 to 9.

Besides the data on the LLs’ participant projects, information regarding the participant teams was also taken into consideration. There were a total of 83 innovators (considering all the members in each participating provider team) that participated in the total six LLs’. Of those 57% were registered as female and 43% as male. The discrimination of the providers’ gender distribution per LL in presented in Figure 3.

**Figure 3.**
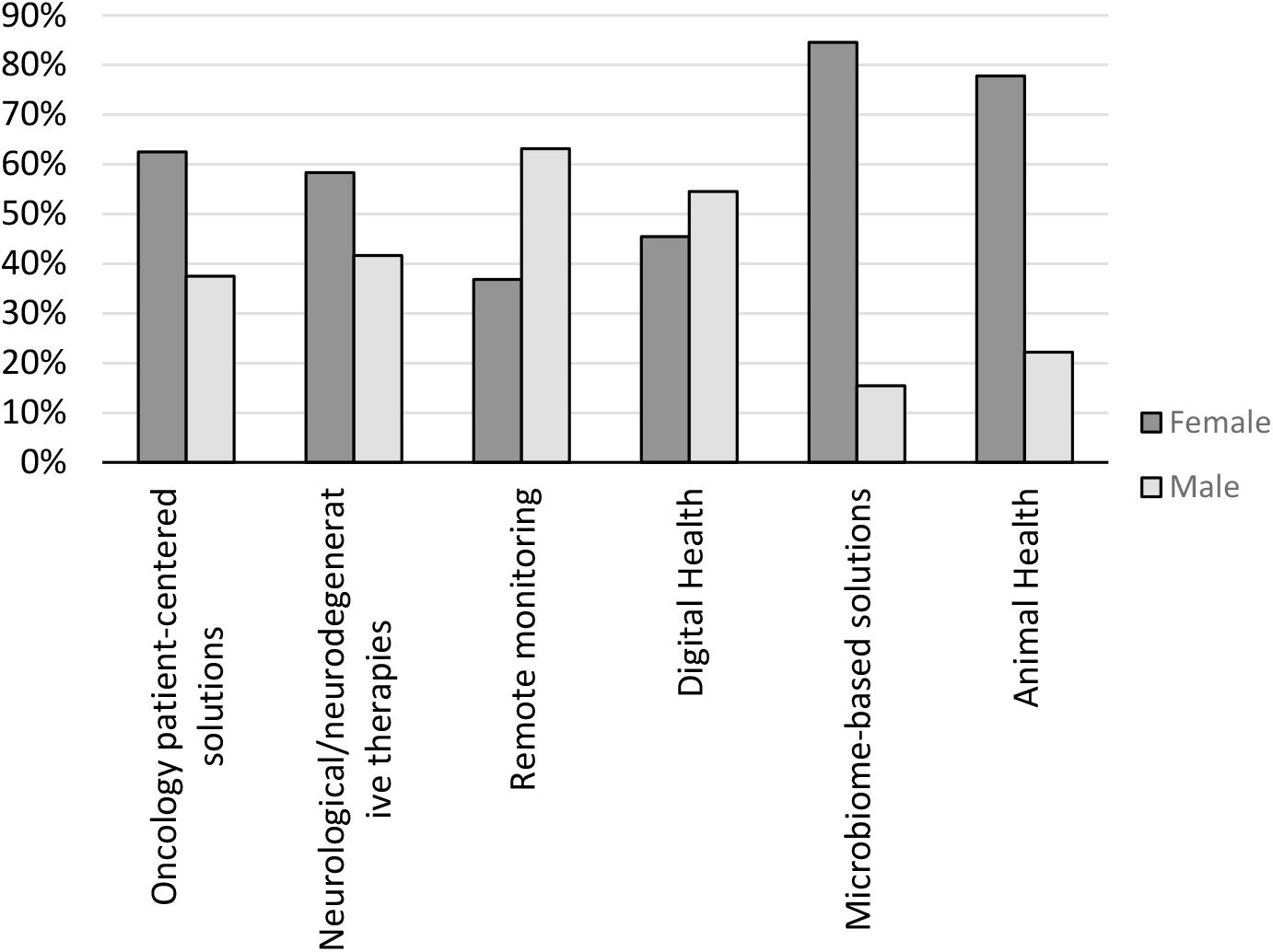
Percentage of female and male innovators that participated in teams per solutions field of application and consequent LL

As it is possible to verify in Figure 3, there were a 67% and 40% higher number of female innovators in the oncology patient-centred solutions domain and neurological/neurodegenerative therapies field, respectively. An even sizeable gender gap was registered in the microbiome-based solutions field and animal health field with 6-fold and 4-fold higher number of females when compared to male innovators, respectively. In contrast, there were a 71% higher number of male than female innovators in the remote monitoring field. A more balanced gender distribution was registered in the digital health innovation domain where only a 20% higher number of male innovators where registered when compared with their counterpart female participants.

### 3.2. Stakeholders

The RESOLVE-HEALTHLLs initiative brought together a total of 77 stakeholders. Of those 54 were end-users and 23 were experts. The number of end-users, specific experts appointed to a particular LL and experts accessed by the providers from all the KIEs, are shown in Table 3.

**Table 3.** Discrimination of the number of end-users, specific experts and general experts per LL.

| LLs | End-users | Experts (Specific) | Experts (general) |
| --- | --- | --- | --- |
| PCCLL | 18 | 6 | 14 |
| TheBrainLL | 15 | 2 |  |
| RemotMonitLab | 13 | 1 |  |
| DigitalHealthLL | 8 | 0 |  |
| MicrobiomeLab | - | - |  |
| AnimalHealthLL | - | - |  |

From Table 3 it is possible to verify that PCCLL had access to 38 stakeholders, TheBrainLL 31 stakeholders, RemotMonitLab to 28 stakeholders and DigitalHealthLL to 22 stakeholders. At this stage, it was not possible to account for MicroBiomeLab and AnimalHealthLL end-user or expert (specific) access. However, had access to the 14 general experts. More insight into the collaborative backgrounds is presented in Figure Data in Figure 4 is discriminated between the end-users and expert stakeholders group.

**Figure 4.**
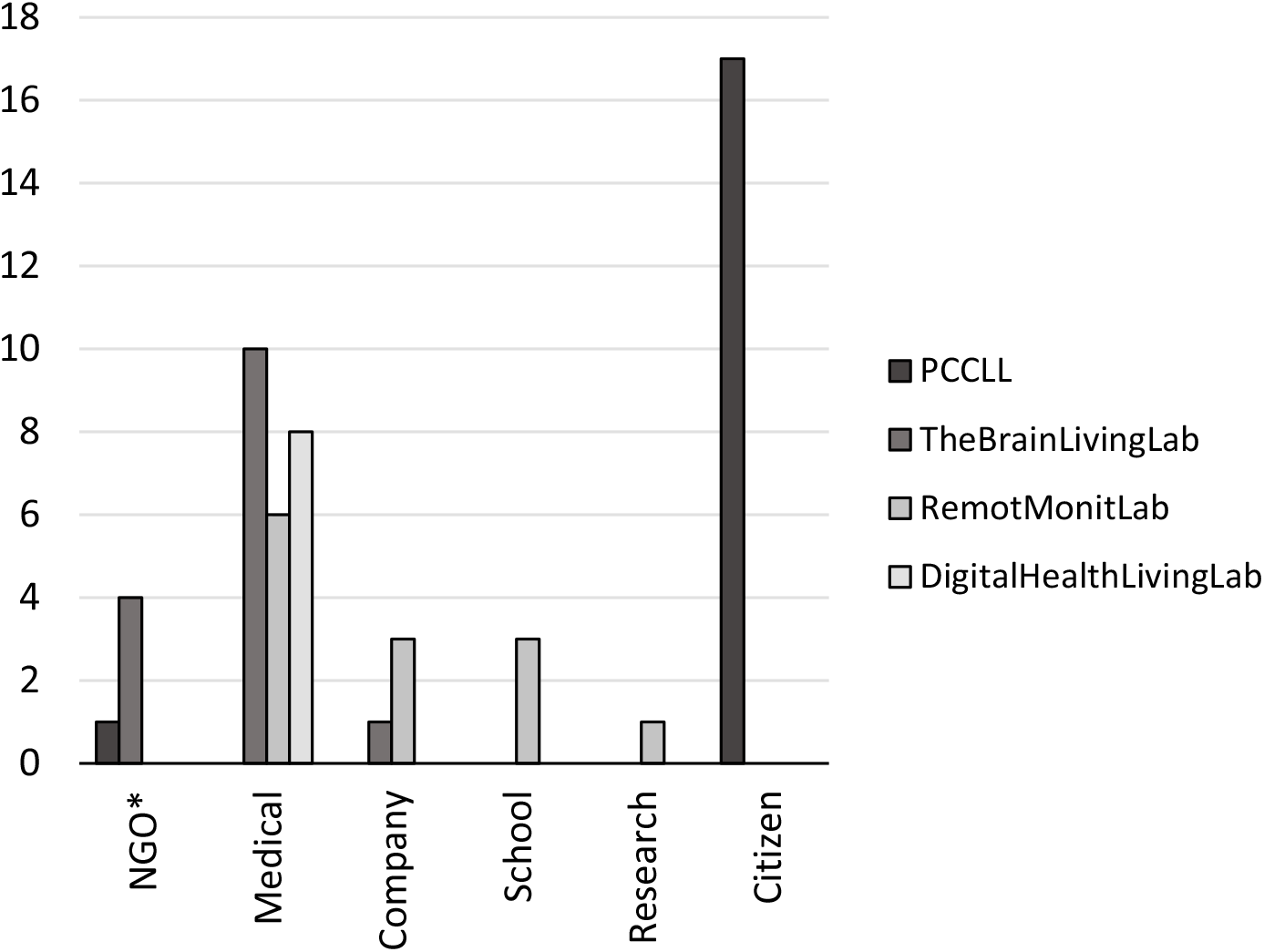
Number of end-users by background type. *non-governmental organization

Figure 4 demonstrates that, except for the PCCLL, for most of the KIEs considered, the majority of the collaborating end-users (44%) were medical system professionals. Although in a reduced number, these LLs also had the collaboration of end-users from non-governmental organizations (NGOs), schools and research. A total of 31% of citizen end-users participated in the PCCLL. In regards to the experts (specialized and general), 61% were from a company background, 22% were from a medical system, 9% were from school, 4% were from a governmental agency and 4% were from research. More detailed information regarding the end-users professional field or background type is presented in Table 4.

**Table 4.** LLs collaborative end-users background.

| LLs | End-users |
| --- | --- |
| PCCLL | Oncological children's parents and personnel from the Associação Acreditar |
| TheBrainLL | Neurologists; neurosurgeons; neuroscience nurses; patient care technicians from the neurology/neurosurgery department; personnel from the Portuguese Association of Parkinson<sup>1</sup> and the Atlantic Association for the Support of Patients with the Machado-Joseph Disease<sup>2</sup> (Psychologist, technical coordinator and social service agent) and translational medicine company manager |
| RemotMonitLab | Dentists; a high school professor and students; a physiotherapist; psychologists; psychiatrist; a neurologist; data science and occupational health physician |
| DigitalHealthLL | Physicians |
| MicrobiomeLab | - |
| VeterinaryLL | - |

<sup>1</sup> in Portuguese *Associação Portuguesa de Doentes de Parkinson (APDPk)*
<sup>2</sup> in Portuguese *Associação Atlântica de Apoio ao Doente de Machado-Joseph (AAADMJ)*

In Table 4 it is possible to verify the diversity of end-users that participated in the KIEs.

### 3.3. Engagement strategies

The LL’s activities were based on a three-stage framework methodology, as illustrated in Figure 5. Each LL adapted and tailored a set of actions in the conceptualization and concretion phase. The actions that took place in the specialization phase were transversal to all LLs and were characterized by the providers from the different KIEs participation together in the same activities.

**Figure 5.**
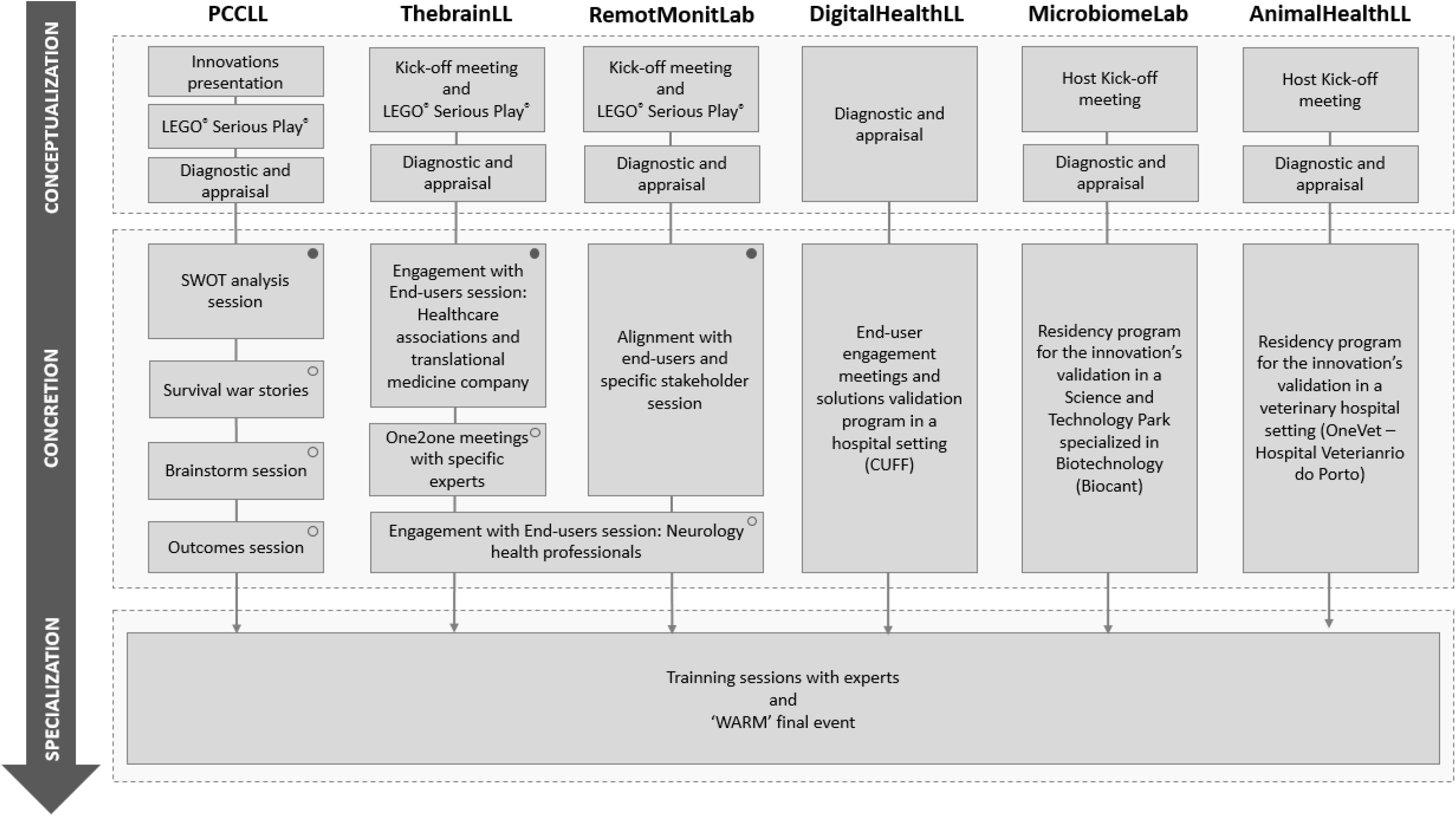
Schematic representation of the actions promoted in each LL with their categorization according to a three-stage configurational process constituted by the conceptualization, concretion and specialization stage. The filled circle are the conceptualization phase’s first session which is developed based on the starting point action and the circle is the following session(s) which are re-designed based on the action research

As demonstrated in Figure 5, in all the LLs, except for the DigitalhealthLL, this phase consisted of two or three activities of ‘familiarization’ with the providers and sometimes also with the end-users and a ‘Diagnostic and appraisal’ session. In the DigitalhealthLL this phase was only constituted by the latter session.

The concretion phase consisted of LL’s specific tailored focus group actions between the providers and the end-users (and specific experts). In the PCCLL, four actions were developed, namely the ‘SWOT analysis session’, ‘Survival war stories‘, ‘Brainstorm session with end-users’ and ‘Outcomes session’. TheBrainLL developed two ‘Engagement with end-users sessions’. The first allowed the involvement of the providers with healthcare associations and a translational medicine company manager and the second with neurology health professionals. The TheBrainLL also provided one-on-one sessions between some providers and specific experts. The RemotMonitLab fostered one action for the alignment between the providers’ end-users and specific stakeholders which occurred in two separate sessions. In DigitialHealthLL the end-user engagement meetings with CUFF health professionals will be provided when required. Also, all the participant solutions will undergo a validation process in the CUFF hospital setting. In both MicrobiomeLab and AnimallHealthLL, providers will undergo a residency program for their innovation’s validation in the Biocant and OneVet real scenario, respectively.

The specialization phase was mainstreamed for all the LLs. Hence, all the KIEs were invited to participate in the following inclusive series of experts interactive training sessions (sometimes with additional one-on-one meetings): ‘IP protection’ given by Anabela Teixeira Carvalho and Tiago Leal from PATENTREE [46]; ‘Health/Medical Software regulatory affairs’ namely ethics committee procedures and data protection requirements given by Miguel Sales Dias from ISCTE-IUL and Cátia Rodrigues and Mariana Abrantes from Complear [47]; ‘Training on data protection in health technologies’ given by Sónia Queiróz Vaz and Joana Mota Agostinho from Cuetrecasas [48]; ‘Legal aspects in the incorporation of companies’ with practices on business and corporate organization namely constitution, governance and forms of financing given by Joana Magina from Cuetrecasas [48]; ‘Supporting tech SME and start-ups’ given by Sofia Bravo from ANI [49], ‘Marketing e branding’ given by Jorge Freitas from Miligram Design [50] and a national innovation incentives session given by Sonia Pinto from IAPMEI [51]. Additionally, interested providers also participated in a ‘bringing with the Investor’, i.e. venture capital (VCs), session which counted with the presence of promoters and investors from Portugal Ventures [52], Faber Ventures [53], Armilar Venture Partners [54] and Beta Capital [55], as well as a Business Angel. During the specialization phase, all the providers were also invited to participate in the health tech acceleration program conference and startup’s exhibition final event named WARM (abbreviation for Worldwide Accelerators Rally at Matosinhos).

### 3.4. Value creation

All the end-users that participated in the PCCLL, TheBrainLL and RemotMonitLab performed advisor, tester and connector roles, with 57%, 38% and 6% contribution respectively, to the ResolveHealth program. The different end-user roles’ percentage contribution per LL is presented in Figure 6.

**Figure 6.**
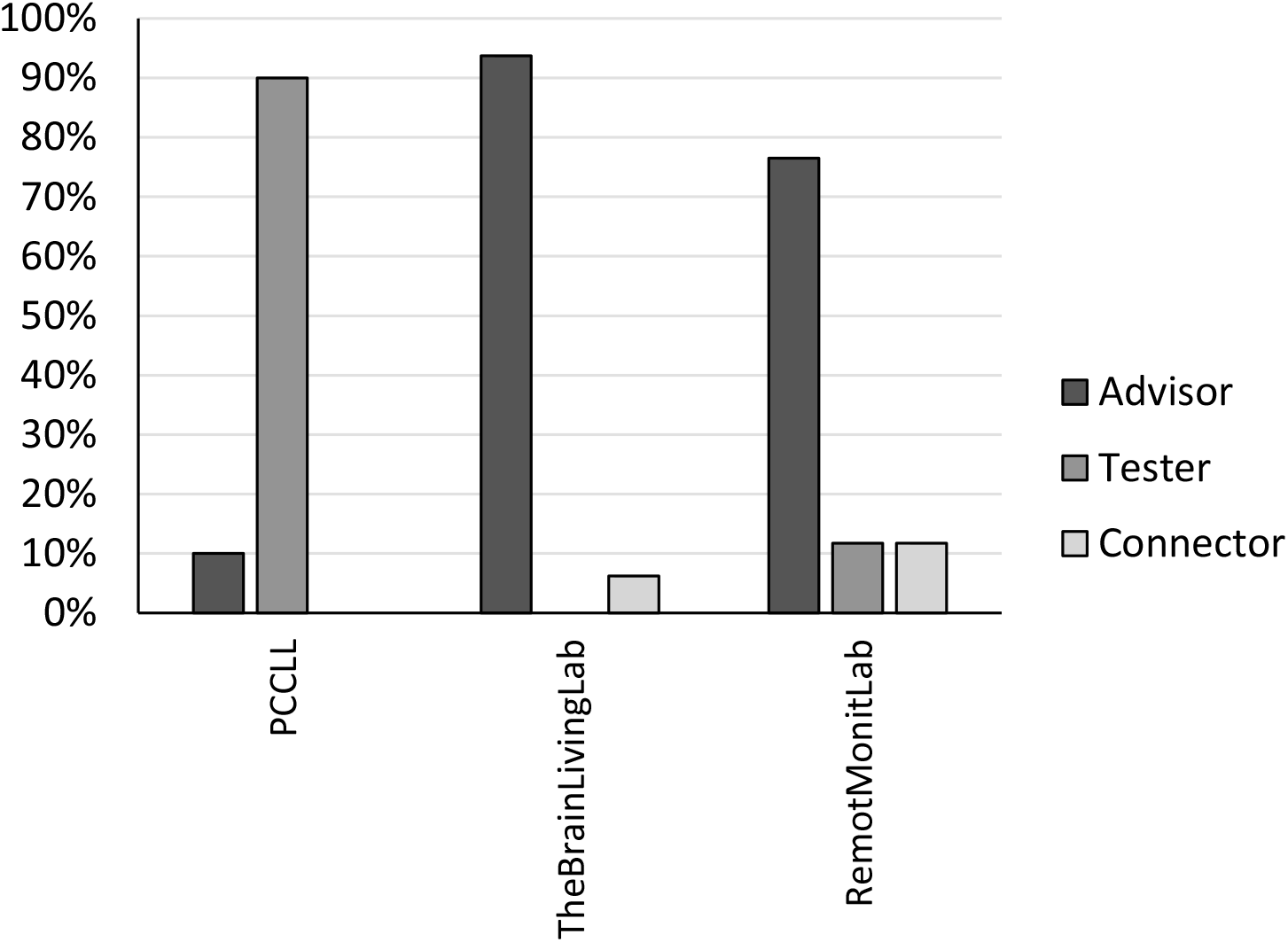
Graphical representation of the end-users percentage role contribution as advisor, tester and/or connector per LL

From the information presented previously and data from Figure 6, it is possible to determine that the advisor is the standard role played by the end-users, which is then followed by the tester role. The end-user role as the connector was infrequent in TheBrainLL and RemotMonitLab and absent in PCCLL. Also, as demonstrated in Figure 6, the RemotMonitLab was the only LL where end-users exercised the advisor, tester and connector roles. The outcomes obtained from the innovations’ participation in the ResolveHealth LLs are presented in Figure 7.

**Figure 7.**
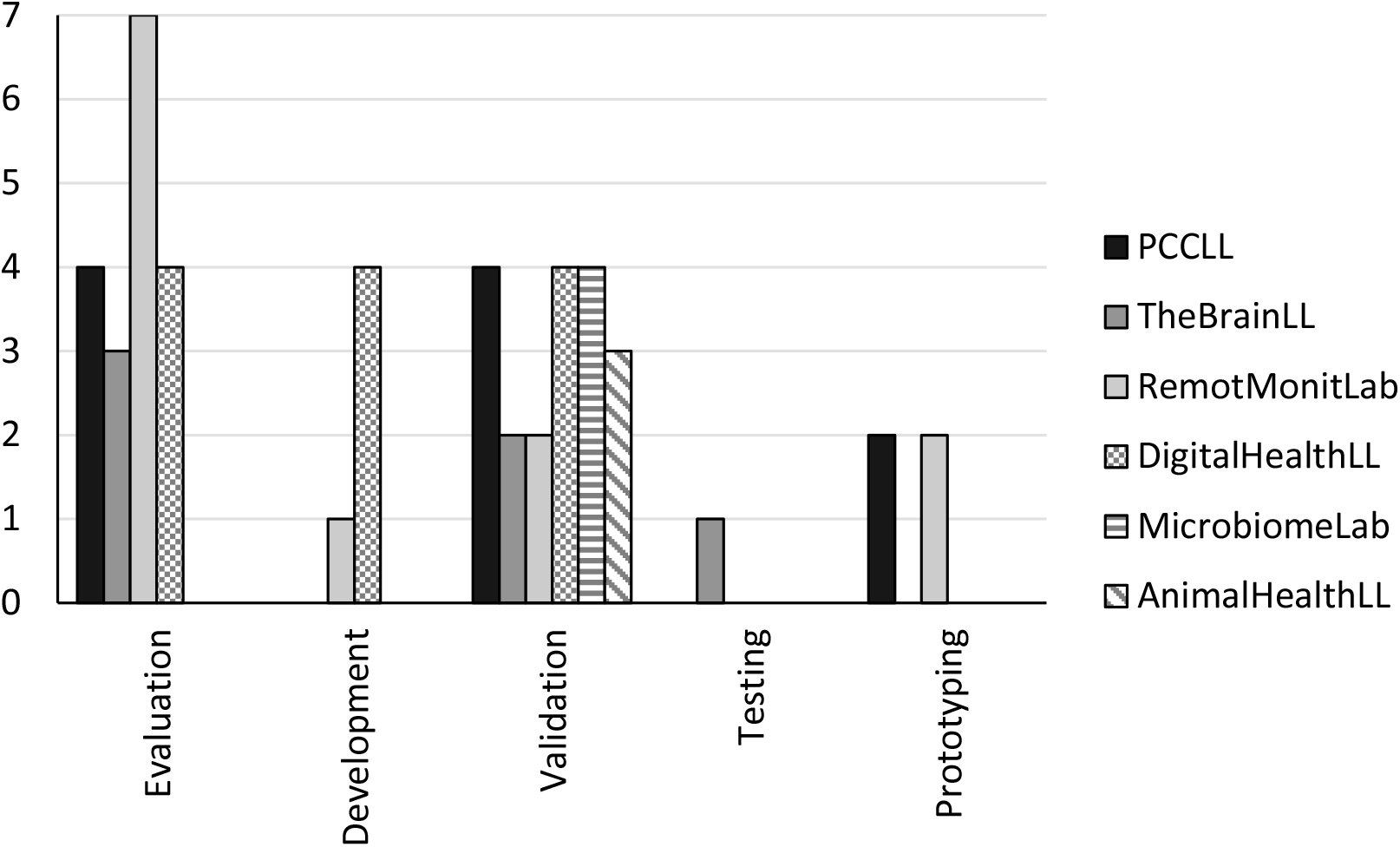
Number of outcomes from their participation in the LL programs

It is possible to verify in Figure 7 that all the LLs had as main outcome innovations’ validation. There is also a high number of innovation evaluations. Both validation and evaluation corresponded to 40.4% and 38.3%, respectively, of the total outcomes obtained from all the LLs considered in this study. This was followed by development outcomes which represented 10.6% of the total outgrowth obtained from participation in the LLs. Exceptional outcomes were the two prototypes development in both PCCLL and RemotMonitLab (8.5% of the total outcomes) and the innovation tested in the TheBrainLL (2.1% of the total outcomes).

Objective feedback on 33% of providers’ participants (each member of the team was invited to participate in the survey) in the LL programs was obtained through their responses to the questionnaire presented in Supplementary Material, Supplementary Figure 2. The results regarding the close answers in the questionnaire are presented in Figure 8.

**Figure 8.**
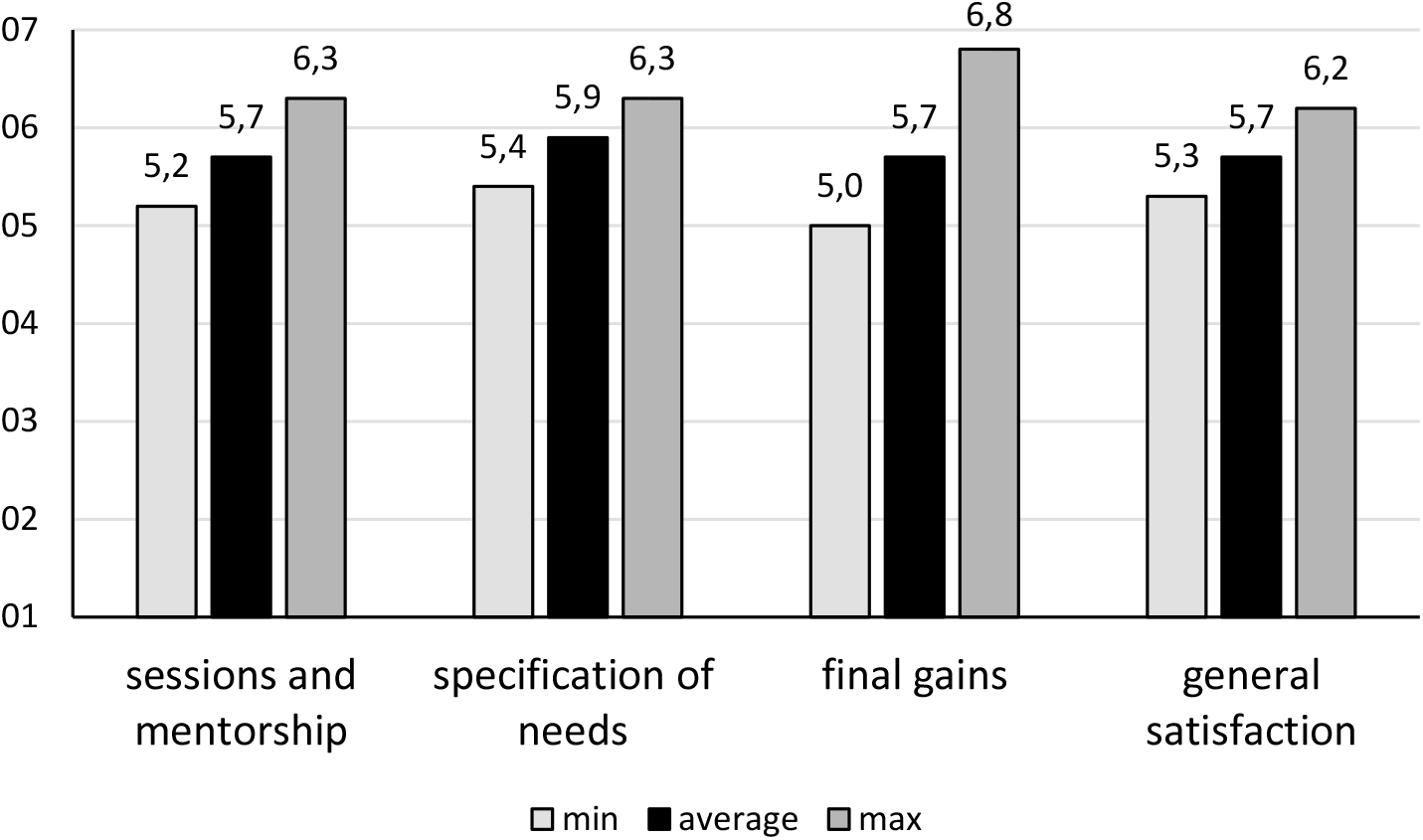
The participant’s response regarding the relevance of sessions and mentorship, specification of actual needs, final gains and perception of general satisfaction experienced during and after they participated in the LL program

As it is possible to verify in Figure 8, on average the LL provider participants demonstrated a high level of satisfaction (between 5.7 to 5.9) regarding the relevance of sessions and mentorship, specification of actual needs, final gains and perception of general satisfaction. Also when considering the standard deviation, the minimum reply value obtained was never below 5. The open surveys allowed more insightful information on the overall program and where revealed providers’ experiences, reactions and concerns regarding the LLs’ activities. The responses obtained from the open surveys are summarized in Table 5.

**Table 5.** Feedback obtained from the survey open question developed to the LLs each provider team member.

| LL | Feedback |
| --- | --- |
| PCCLL | "This is a highly relevant leverage program. I highlight four aspects: 1) being under the hat of i3s and being managed by staff from this institution; 2) finance factor was a game changer in the support landscape of national entrepreneurship; 3) complementary training via workshops/seminars and contact with stakeholders and 4) access to end-users to be accompanied by program mentors. What would I add? One-on-one interactions with mentors from companies aligned with the activity of the projects (non-competing), to discuss corkscrew tactics and solutions (so useful in the nascent phase of projects trying to access the market). Super grateful for the professionalism, dedication and fairness of the entire Resolve health team." |
| TheBrainLL | - |
| RemotMonitLab | "The speakers were relevant and knowledgeable, and the sessions were interesting. It was useful to have the sessions in an online format"<br>"The program was very well organized. The only issue was that the program for the different sessions was not very clear when the program started. A brief |
|  | meeting with the participants explaining the timing and mentoring process would have been very useful.”<br>“Unfortunately, our expectations for the program were not met at all. The program's scope was conducting a pilot, and although the initial meetings with the Resolve-Health team pointed in a good direction for that, the development of the conversations and process with the entity that we selected for the pilot, together with the Resolve-Health team, were not successful. After the program has already finished, we still haven't a final response from i3S. Although we understand that the internal processes require the approval of the Ethics Committee, we believe that on these types of testbed programs, the team should have a defined protocol to target the companies/organizations for pilots and be better informed about the internal processes they require. Furthermore, as we mentioned to the team several times, our expectation regarding the financial support was also not met, given that the regulation of the program stated that each team could ask for up to 20K€ of support, given the needs. We explained that we would need a certain amount (around 8K€) to develop a new feature to our product, which was also in the scope of the program, but we ended up only receiving 3K€ and no pilot. We hope that this feedback helps the team improve the program, and we appreciate the efforts made to satisfy our needs in the best way”<br>“I would suggest that all sessions can be assisted via Zoom. Some people live on the Islands, which makes it difficult to travel. Other than that, it was really well organized and we learned a lot.” |

As it is possible to verify from Table 5 there was only a reduced feedback regarding the provider’s participant’s experiences, reactions and concerns. There were no observations from the TheBrainLL provider team members. The enabler’s feedback on their participation in the LL is summarized in Table 6.

**Table 6.** Feedback obtained from e-mail contact with each LL’s host manager.

| Enabler (LL) | Feedback |
| --- | --- |
| Ana Monteiro from Associação Acreditar (PCCLL) | *“I take the opportunity to underline that I really enjoyed the partnership with you! It was very productive for the families, they gave us very interesting clues and the articulation with you was great. Congratulations for your work”* |
| Jorge Alves from Centro Cerebro (TheBrainLL) | *“Congratulations on the organization of the WARM event”;* |
| Marques Teixeira from Neurobios (RemoteMonitLab) | - |

### 3.5. Results dissemination

LinkedIn and the Resolve-Health website were the main social media considered for the promotion and dissemination of the LL’s actions and outcomes. LinkedIn had 1345 followers on July 30^th^ 2023 data collection. From those, more than half of the total followers are from the Research field (25.1%), Education (14.3%), Business Development (7.7%) and Healthcare Services (7.1%). There was a total of 11.7% international followers. In regards to the demographic location of the national followers, 44.4% were from Porto, 17.2% from Lisbon and 16.3% from other Portuguese main cities (namely Braga, Coimbra, Guimarães and Aveiro). The number of articles on LinkedIn with information regarding the actions exclusively developed during the conceptualization and concretion phase by each LL was: 5 for TheBrainLL, 4 for each PCCLL and RemotMonitLab, 3 for each and DigitalHealthLL and MicorBiomeLab and 2 for AnimalHealthLL. There were a total of 19 posts published on LinkedIn related to the LL’s specialization phase activities. The average engagement rate and standard deviation generated in LinkedIn by the totality of posts published in each LL and regarding only the activities that were developed during the conceptualization and concretion phase as well as those published during the specialization phase are presented in Figure 9.

**Figure 9.**
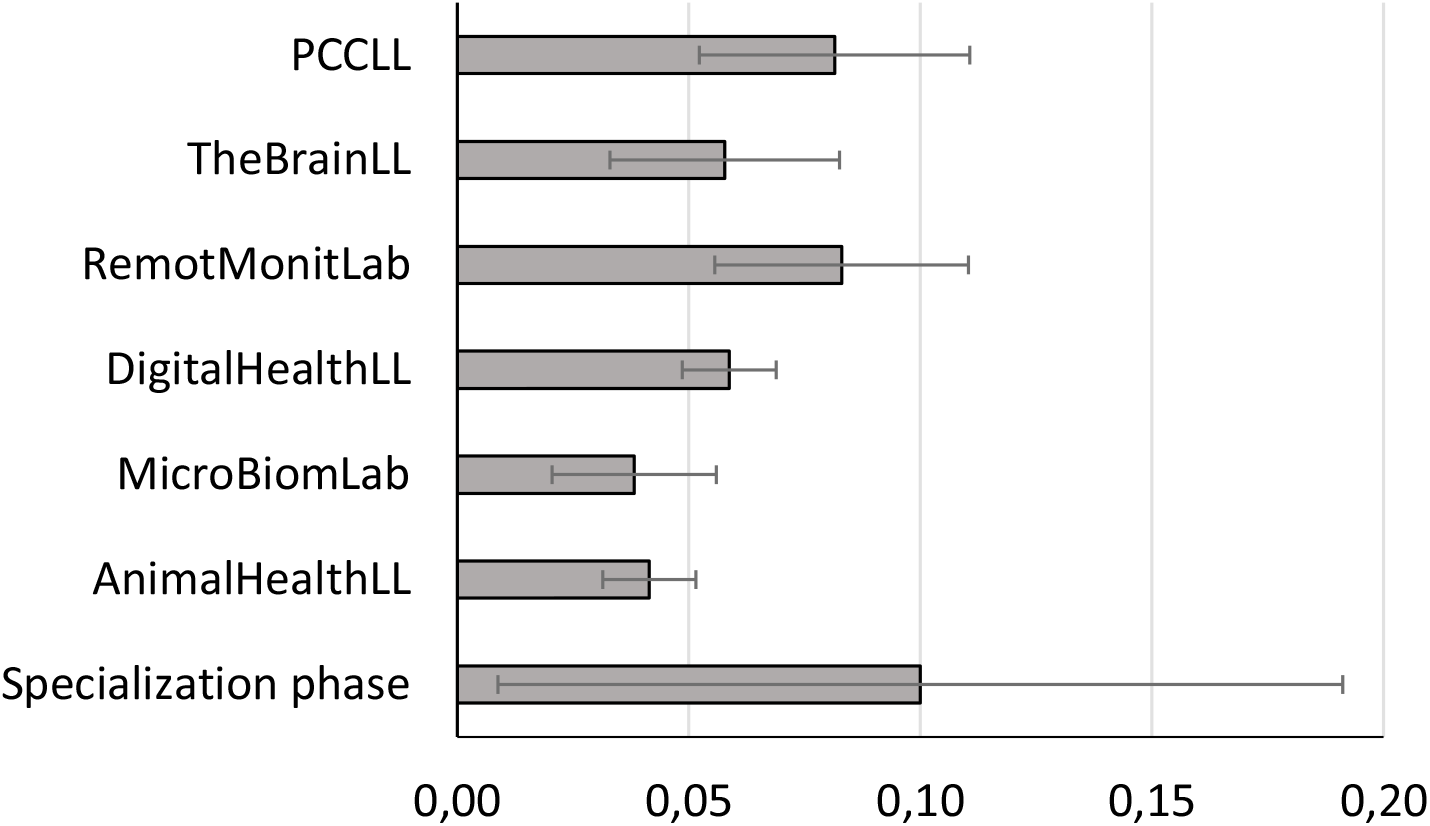
Linkedin average and standard deviation engagement from the totality of the posts published regarding the conceptualization and concretion phase activities per LL and from the posts published about the activities developed in the specialization phase

From Figure 9 it is possible to verify that both PCCLL and RemotMointLab generated the highest engagement. The MicroBiomeLab and AnimalHealthLL were the KIEs which attracted less public attention by a generation with the lowest average engagement. It is also possible to notice that although the specialization phase posts generated the highest average engagement rate they were also associated with the highest standard deviation value. From a more general perspective, a total of 606 unique page visitors were registered. Of the total page visitors 25.4% were from the Research Services Industry, 20.8% from Hospital and Health Care, 10.2% from the higher education field, 7.5% from the Biotechnology Research domain and 4.2 from the field of IT Service and IT Consulting. Figure 10 represents the engagement rate percentage between November 2022 and July 2023 as a result of the publication of posts that contain exclusive information related to the Resolve-Health program LL’s activities.

**Figure 10.**
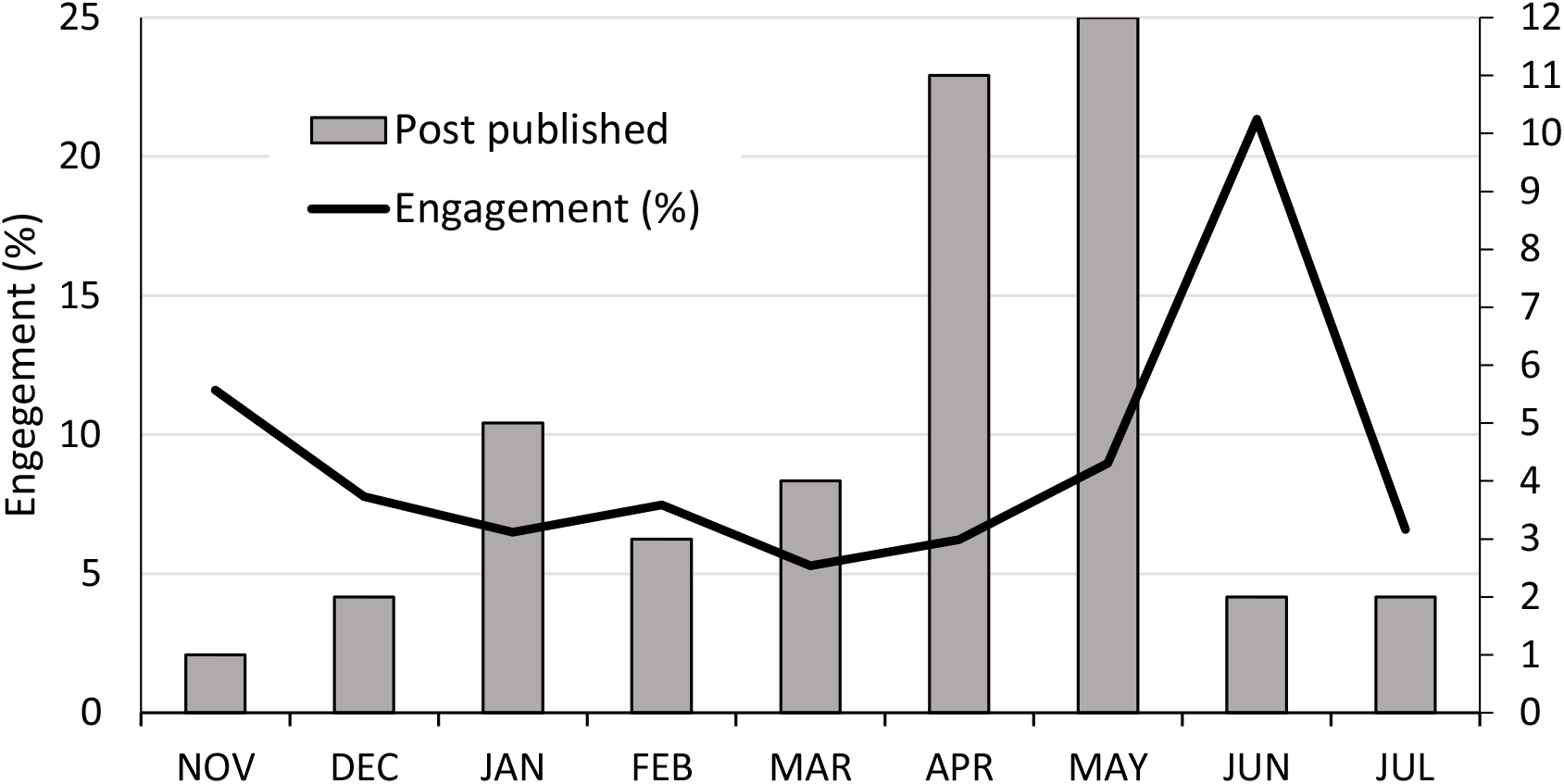
Number of posts (grey bars) for all the LL’s activities published and the average percentage engagement (black line) per month from November 2022 to July 2023.

It is possible to verify from Figure 10 that from November 2022 to July 2023 the average engagement percentage was considerably constant except for June. Also from this Figure, it is possible to notice that April and May had a particularly high number of posts published (11 and 12, respectively) when compared to the other month’s average of 3 posts per month. The Resolve-Health website was used for the promotion of the LL open calls and the presentation of the teams supported by the ResolveHealth2.0 KIE program. The publications developed per LL are summarized in Figure 11.

**Figure 11.**
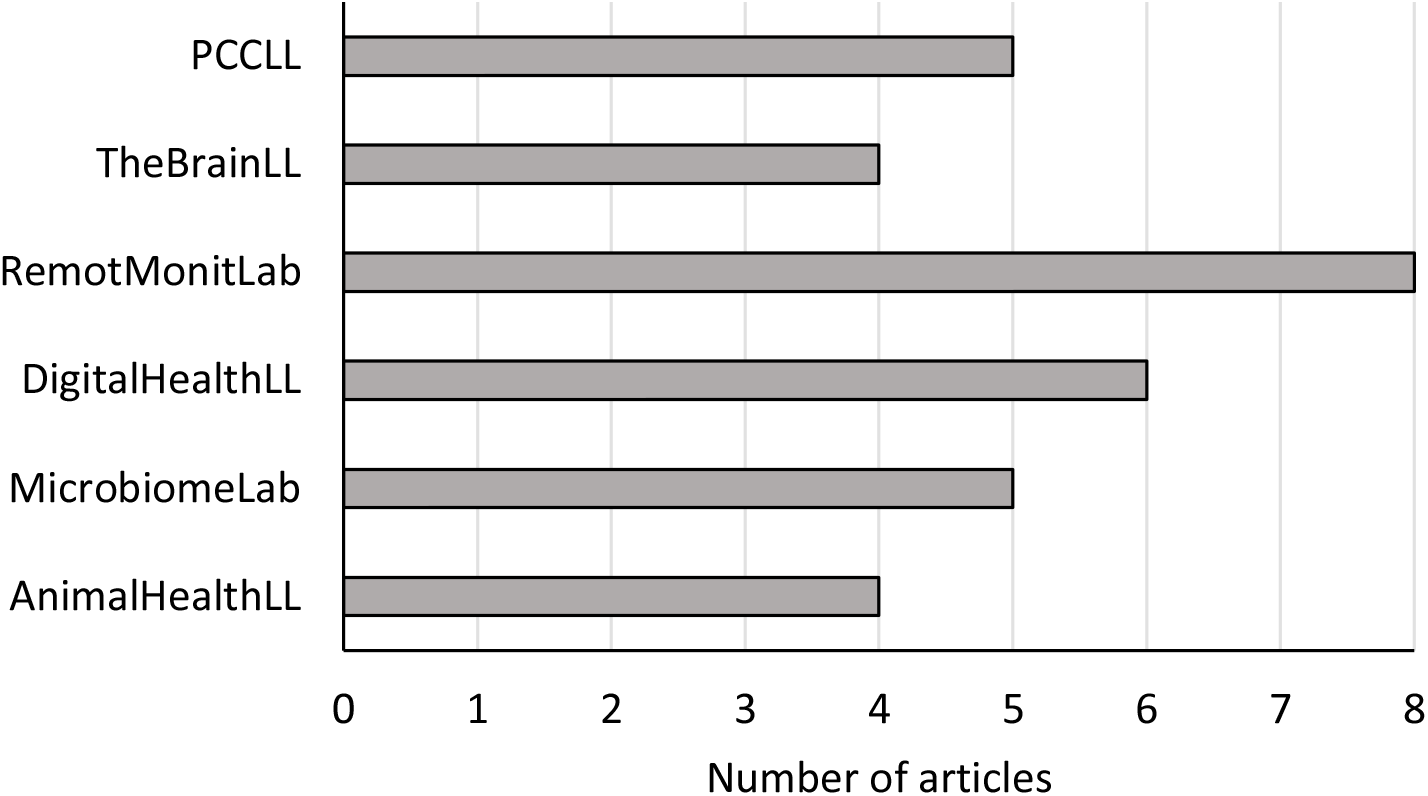
Number of articles per LL posted on the ResolveHealth official website

From Figure 11 it is possible to verify that RemotMonitLab presented a significantly higher number of articles published. TheBrainLL and AnimalhealthLL were the KIEs that published a reduced number of articles during the ResolveHealth program. No information regarding the engagement with the webpage was obtained.

## 4. Discussion

The Resolve-Health program LLs’ presented in this article were from different application contexts within the health domain which allow for the discussion and proposal of a set of efficient and standardized good practices to be applied in similar KIEs. The ten good practices found from the singular and cross-case analysis of the six LLs were: strong marketing and promotion strategies of LLs’ open calls; enforcement of gender equality; proper stakeholders’ selection; development and implementation of suitable framework strategies for end-user engagement and value creation; value creation for all the LL partners; lengthening of the long-term viability of the LL project; LL actions and impactful promotion and dissemination of results; fomentation of environmental sustainability and results information processing for LL performance evaluation.

### 4.1. Marketing and promotion strategies of LLs open calls: The determinant step for ensuring that the LL tool reaches the appropriate innovators

Any promotion and marketing need to be based on a strong communication strategy using a variety of digital platforms and other communication methods in a sustainable way [56]. With this in mind, aligned marketing and promotional strategies for the advertising of the open calls were adopted for all the LLs’ considered in this study. Accordingly, in an attempt to increase the level of awareness of the project [56] a graphic designer and a communication manager were both an integral part of the Resolve-Health program team. Hence, communication actions through social media (*i.e.* LinkedIn, Facebook and Instagram) were harmonized and integrated into a uniform defined and easily recognizable branding and design. For the advertising of the six LLs’ open calls, as described in the methods section and presented in Table 1, each LL call provided information regarding their areas of action and the type of innovations supported (*e.g.* new service, new product, new therapy etc.). A rigorous clarification of the support provided for each LL program was also included in the promotion strategies, as suggested by Alavi *et al.* [57]. As stated by the author such an explanation is essential to guarantee the candidates’ understanding of what outcomes are expected from the KIE approach and that would not have been achieved through less demanding methods. Ignasi Capdvilla [58], however, accentuates the importance of clarifying the methodology adopted by the LL since there are no standardized defined approaches and therefore multiple practices are adopted within the KEIs. This information was not provided in detail in the LLs’ promotion strategies since in this study the KIEs were developed based on an AR methodology (this topic will be discussed in detail in section 4.4). Hence, the methodology was adapted or created based on a *feedback* approach to actively enhance the engagement between providers and stakeholders for the improvement, and validation of groundbreaking products, services and technologies.

Although similar open-call marketing and promotional strategies were adopted for the six LLs, different advertising times were required to attain the minimum number of project applications. As indicated in Table 1, the open-call duration for LLs considered in this study was on average 3 weeks. The AnimalHealthLL, RemotMonitLab and DigitalHealthLL showed the lowest required promotional time (2 weeks). The values obtained for RemotMonitLab and DigitalHealthLL could most probably be justified by Irma Mäkäräinen-Suni [12] statement that ‘new technology as a theme for Living Lab is practically marketing itself’. The AnimalHealthLL was expected to be launched within a tighter schedule which limited the promotional time to an established 2-week maximum time and not the number of applicants threshold. Despite the general marketing hype associated with LLs, PCCLL and TheBrainLL, required a prolonged promotional time of 6 and 5 weeks, respectively, to obtain a considerable amount of applicant candidates. It was possible to assume that this phenomenon was caused by the narrowing of these KIE specialization fields which reduced the available innovation projects to a niche segment. We also found that the KIEs programs’ economic support announced brought about a considerable increase in the number of applicant projects. The “enthusiasm” for the economic support was verified during the interviews where several candidates justified this feature as the incentive for the application of their project to the LL. As stated by a PCCLL provider the “finance factor was a game changer in the support landscape of national entrepreneurship” (see Table 5). However, it is important to highlight that this capital translated into different forms of financial support such as money, coaching, mentoring, consulting services and infrastructures as it is possible to see in Table 2.

The LL area of concerned not only influenced the advertising time but also reflected in the number and type of participant innovations. Hence, whereas the PCCL specialized in supporting e-health solutions for oncological patients’ caregivers, DigitalHealthLL had a wider area of action which includes digital technologies to improve health and healthcare. The openness to new technologies for the general healthcare application is reflected in a 4% higher number of participant projects in the DigitalHealthLL when compared to PCCL. In the TheBrainLL and RemotMonitLab LLs, the areas of action are closely related being both in the neurological field and in some cases the supported innovations might even have a complementary role. However, while TheBrainLL focused solely on therapies for neurological disorders, the RemotMonitLab LL specialized in technology-based solutions for mental health support. Hence, there was a wider range of innovations which could fit the requirements for remote monitoring technology instead of therapies led to a 15% increase in the number of participant projects in RemotMonitLab when compared to the TheBrainLL. We might also assume that the distinct number of projects that participated in the RemotMonitLab (when compared with the other LLs) could be an outcome of the market increase demand and accelerated propensity for remote healthcare delivery technologies as a consequence of the stay-at-home and social-distancing rules implemented during the COVID-19 pandemic [59]. This assumption can be supported by Tang *et al.* [60], which stated that remote patient monitoring in traditional medicare increased substantially during the COVID-19 pandemic, reaching more than 6 times the pre-pandemic levels by September 2021. The high number of participant projects for both the RemotMonitLab and DigitalHealthLL when compared with the other considered LLs reflect the market focus on advancement trends in the creation and integration of smart healthcare technologies [61]. The accession to the development of these technologies is intensified by the firm basis for further development provided by the Internet of Things (IoT) technology [62], and artificial intelligence (AI) [63] in supporting person-centred care. The AnimalHealthLL distinguishes itself by being the only LL focused on animal health against the other LLs which target human health solutions. This is reflected in a lower number of projects that participated in this LL. Although it is not easy to justify the reduced adhesion to this LL, it could be explained by 1) this LL’s limited promotion time and 2) the underestimation of animal health solutions’ impact on the global economy, human health and sustainable development despite animal disease and health challenges constant evolvement [64]. The MicrobiomeLab also has a distinct action field by supporting microbiome-based solutions. Also as shown in Table 2 the participant innovations within each LL varied considerably in terms of technology/service provided, TRL, specific policies adopted (*e.g.* IP protection strategy) and technical support provided. Besides TheBrainLL in which participant projects showed similar TRL levels, there was a considerably variation in the TRL of the solutions that participated in the KIEs. It was also very interesting to verify that, regardless of the LL considered there was a diversified range of promising new solutions which addressed complex market challenges.

While there is no standard framework for LL open call promotion, it is clear that the digital tools considered were a decisive factor for reaching out not only to innovators and entrepreneurship parties form different organizational groups such as universities, R&D institutions/centres, spin-offs, start-ups but also to the wider public. By adopting this strategy it was possible to promote the open calls at a local and regional level and also within and outside the national territory. Despite the Resolve-Health LLs program’s openness to supporting innovation projects at an international level (*e.g.* Luckietech and Virtuleap), the majority of the participant innovations were from the national territory. The catering for national innovation projects allowed us to infer about the Portuguese innovation ecosystem. Notwithstanding the significance of the promotional tools considered in this study *i.e.* Resolve-Health website and social media (namely LinkedIn, Facebook, Instagram and twitter) we would like to highlight other impactful marketing media suggested in literature which could amplify the call for the appropriate LL participants accession such as blogs [56], email newsletter [65], press release through local and or national newspapers [66, 67] and radio announcement [66].

#### Open call promotion plays a crucial role in ensuring that the LL tool reaches suitable innovators

This step has a considerable impact on the KIE course of action since the participating projects, will inadvertently, condition and rule the LL’s future activities. Hence, the impact of the LLs’ open calls marketing and promotion strategies as an essential good practice was highlighted in this section as well as its influence on the number and diversity of projects applied.

### 4.2. Gender equality: Fight for gender equality and the empowerment of women

The United Nations (UN) Member States Sustainable Development Agenda (Goal 5) aims to achieve gender equality and the empowerment of all women and girls by 2030 [68]. This concern was shared by the Resolve-Health program, whose core team presents an equal gender distribution and contribution to the development of the actions. Additionally, the criteria for human resources recruiting and selecting was based on the principle of meritocracy as well as on the principle of equality and non-compliance which guaranteed no discrimination based on gender. Conditions were also disclosed for the insertion and development of skills on equal terms for men and women, guaranteeing a safe, healthy and non-discrimination work environment. For this purpose, the Resolve-Health program adhered to the European Commission initiative described in the "European Charter for Researchers" and in the "Code of Conduct for the Recruitment of Researchers". Despite, the teams’ equal gender distribution or the presence of female leadership in the projects not being considered a favouritism criterion for the innovations selection, there was a considerably overall balanced innovators gender distribution in the LL with only 14% more female than male innovators. Hence, the resultant participant projects team members’ gender percentage distribution corresponds to an un-bias realist’s representation of the innovator’s ecosystem in the studied LLs. However, when analyzing LLs individually it was possible to verify this balanced tendency was only registered in the DigitalHealthLL where there is only a 20% higher number of male than female innovators. This favourably contradicts literature data [69] that stated the existence of disparity between male and female representation in the development of digital health innovation. A female dominant representation was present in PCCLL and TheBrainLL with an even more pronounced gender disparity registered in the MicrobiomeLab and AnimalHealthLL. The sharp tendency registered in the MicrobiomeLab (6-fold higher number of female innovators when compared with the male counterparts) is in agreement with recent data published by the European Patent Office (EPO) which rated biotechnology as the field with more women inventors [70]. The female inventor’s dominant representation in the LLs could most probably be a reflection of the Portuguese higher number of female researchers in the public sector (57,6%) [71, 72] and their also higher representation in the medical and health science field. According to statistical data [72] in Portugal women are the majority in virtually all research domains, except in Exact Sciences, with only 32.8% of women and in Engineering Sciences and Technologies, where the feminization rate is only 27.7%. In contrast, there was a 71% higher number of male innovators when compared with the female ones in the RemotMonitLab. The male innovators’ over-representation can be explained by the consistent gender gap present in the STEM (science, technology, engineering, mathematics) field [73, 74]. In both the EU27 and Portugal, the number of men who graduated in these areas is always significantly higher than that of women [72]. As stated by Ahmadi *et al.* [75] in the IT sector, participation is still unequally distributed with female programmers, software developers and engineers being a minority. The authors also stated that factors like cultural stereotypes, hostile environments, gender wage gap, a lack of advancement or missing role models and peers among others are reasons for women showing less interest in this field. Despite the unsettling disparities registered in the STEM field, women are in general less visible as innovators than men even though there are strong indicators that gender composition may affect a firm’s innovation capacity [76]. According to a study developed by the EPO [70], when considering data from 1978 to 2019, only 3.2% of inventors in Europe are women and although the women inventor rate in Europe has been rising in recent decades (up from only 2% in the late 1970s to 13.2% in 2019) a strong gender gap remains. Although there is no clear explanation for these differences, since gender has not yet been extensively explored in innovation studies [71], it is believed the fact that the problem-solving skills of women in male-dominated sectors are less likely to be regarded as an innovation should be taken into consideration [71, 77]. It is not only in innovation development that women face gender-related challenges since they continue to be confronted with numerous setbacks in starting and expanding their businesses. Since, limited access to markets, technology, networks, and financing are determinant features in their involvement in entrepreneurial activity [78]. Raman *et al.* [79] also suggested that factors such as women’s lower self-perception of their skills when compared with their male counterparts may prevent them from engaging in entrepreneurship or applying for finance. So even when women embrace the entrepreneurship path, according to research developed by Brush [80], they receive less than half the level of investment than their male counterparts. An underlying factor which is believed to influence this phenomenon is the fact that 93% of VCs are men who thoughtlessly or not are more likely to invest in male entrepreneurs [81]. Consequently, discrimination based on gender has a significant impact on how entrepreneurial ecosystems are developed. Hence, this raises the question of how the KIE can promote gender equality in innovation and women entrepreneurship in the health-related field but mainly in the STEM health innovation sector. As an example there is the inspirational project developed by Ahmadi *et* al. [75, 82] where LL was considered in the context gender and IT and as a tool to reveal new and relevant insights and create social change in a collaborative way. Although there is an emergence of novel KIEs initiatives for developing and supporting solutions primarily aimed at addressing problems faced by women [83] and/or to promote gender equality [84, 85], both approaches still remains under-explored.

There is an urgency to avoid the loss of competitive market innovation potential as a result of women’s lower propensity to actively pursue efforts to develop knowledge, innovation and entrepreneurship, especially in STEM-oriented fields of study. Hence, it is important to develop and design the KIEs based on frameworks that promote research developed by gender inclusiveness teams and that provide an ecosystem that attracts female researchers and innovators and helps to reduce the imbalances found. Possible interesting actions would be to in the scope of the LL 1) raise awareness of the existence of a gender gap in innovation and entrepreneurship, 2) increase the participation of the under-represented gender by favouring the selection of female-led innovations projects in specific fields where there is a marked gender gap, 3) monitor the participant female innovator interest and motivation in the LL activities, 4) develop brainstorming sessions on how to adapt the LL activities to better support female entrepreneurship, 5) develop lectures to highlighting opportunities for female leadership, 6) the sharing of success stories to break the prejudice associated with female entrepreneurship and 7) support transdisciplinary innovation which is also associated to enhance gender equality projects. We would also like to highlight the urgency of not only **fighting for gender equality and the empowerment of women** in the innovation and entrepreneurship ecosystems but also the importance of considering the integration of innovators from social categories intersecting with gender such as ethnicity, disability, sexual orientation or else social origin into the KIEs.

### 4.3. Stakeholders’ selection: Without their participation, the LL will fail

It has been long established that the physical and social context in which an activity takes place is likely to be relevant to how that activity unfolds [57]. Hence, a strategy based on a thorough analysis of the promoter’s expectations, innovation needs and specificities while placing them in the respective KIE area of application was the driving force for gathering a set of suitable stakeholders for the LLs sessions. Also taken into consideration was the stakeholder’s background since they will not only share personal ideas and inputs and understand and evaluate technology use within specific situations but will allow the providers to extrapolate the information provided to a wider population they end up representing [14]. Consequently, previous successful and trustworthy collaborations with the Resolve team and previous collaborations with the host’s team members were the starting point contact for the gathering of the most suitable stakeholders. Since their interests and motivations will influence their interaction and contribution to the LLs [1], stakeholders’ initial motivation level was also a parameter taken into consideration during their selection phase. In the Resolve-HealthLL program, there was a distinct role played by the end-user and expert. This led to the gathering of specific individuals as end-users who would be able to contribute to the participant innovations leveraging through “co-creating”, testing, validating and/or development. On the other hand, the experts were agents who provided specific knowledge on a particularly complex subject (*e.g.* commercialization or marketing) where there is little information or even a considerable degree of uncertainty [86]. Particularly given our experience, and as emphasized in the literature [1], it was easy to captivate the attention and interest of both end-users and experts to collaborate in the LL program. We would like to highlight the paramount role of the LLs’ liaisons whose responsibilities varied from public relations, organizing the KIE and finding partners (as a stakeholder mobilizing agent) but also, as stated by Welling *et al.* [56], as relationships builders with the different parties and creators of a positive reputation for the LL and within the stakeholders’ communities. In these particular LLs cases the liaisons also as the additional role of a research study conductor. Contrarily to what was suggested in the literature [1, 57], when evaluating the end-users’ backgrounds (Figure 5 and Table 4), it was not possible to guarantee, for all the KIEs, that there was one representative from each background classification sector (Government, NGO, Medical, company, school, research and citizen). Although, citizen end-users are seen as valid and important providers and producers of knowledge that shape innovation [87], in both TheBrainLL and RemoteMointLab there was no indication that there would be an additional benefit in having them present during the LL program. For example, in TheBrainLL the presence of patient end-users (*i.e*. citizen end-users) was purposely avoided to reduce the expectations from the presentation of pharmacological therapies for the recovery or management of incurable and/or rare diseases (*e.g.* Machado-Joseph disease). The same caution was adopted in the RemoteMointLab where technology-based solutions for mental health problems were being supported. We would like to point out that, albeit the NGO type was considered in this study, we do not believe that this category would be a “must” in all health-related KIEs. Hence, the NGO can be considered as a facultative or complementary category of end-user.

We would like to highlight the importance of having end-users from a health-related government background since they can provide economic and human resources, as well as, physical structures (offices, meeting spaces and laboratories) and additional opportunities for national and international collaborations due to their extended networking [1]. However, it is challenging to have these end users present in the LL’s activities since they are normally unavailable or less readily available. This challenge needs new strategies to be defined.

An interesting government end-user partner would have been for example the collaboration with the TheBrainLL of an agent from Infarmed [88] which is the Government agency accountable to the Portuguese Health Ministry that evaluates, authorizes, regulates and controls human medicines as well as health products, namely, medical devices and cosmetics for the protection of Public Health.

The selection strategy adopted allowed for a considerable overall number of collaborating stakeholders to participate in the LL program. Of those 53 were end-users and 23 experts (Table 3). However, this value varied between KIEs. As it is possible to infer from Table 3, the number of end-users per number of participant innovations rate was 5 for TheBrainLL, 4.5 for PCCLL and 1.9 for RemotMonitLab. Although data for the DigitalHealthLL was presented for reference purposes in the results section, it will not be considered subject to comparison since this LL program is still in its early stages. It was also interesting to verify that besides PCCLL, which developed a centralized overview focused mainly on a specific group of final end-users, both TheBrainLL and RemotMonitLab foster the participation of end-users from a wide variety of backgrounds within the healthcare domain. As it is possible to verify from Table 4, the main end-users in the TheBrainLL are neurology medical field-related health professionals and personnel from neurodegenerative diseases associations, as well as, a highly regarded translational medicine company manager. These end-users are ultimately, those who could guide the teams towards the development of innovations with strong market application and based on realistic expectations. The RemotMonitLab showed the broader diversity in end-users backgrounds, from dentists to a high school professor and students, a physiotherapist, psychologists, a psychiatrist, a neurologist, a data science professional and an occupational health physician. This variability can be justified by the attempt to provide suitable guidance and collaborations to an also considerably diverse set of participant innovations. According to Anna Ståhlbröst [3], this variability improves the quality of the LL services being developed. Since knowledge, both scientific, technological and sector-specific, is considered a critical innovation factor [89], the multiple perspectives increase its collective sharing as well as enhance creativity which brings power to the development process and contributes to the achievement of rapid progress [3]. This was further evidenced in the engagement with end-user LL sessions where the number of collaborators, but more importantly the heterogeneity in their backgrounds (*i.e.* diversity in the expertise, life experience and knowledge present) was more notable. This diversity contributed to immersive dialogues, multi-perspective debates and interesting reflections between the end-users and between the end-users and providers. As a result, during those sessions, there was an enhancement in the quality of the inferences obtained and feedback provided. Despite the variety of end-users present in the KIE sessions, the collaborations developed were in most cases the result of two-way or three-way relationships.

By allowing the providers and their solutions to contact and engage with both end-users and experts it is possible to synchronize and nurture the process of knowledge generation and innovation progress from a quadruple helix model perspective [1, 90]. As stated by Nguyen and Marques [91] by involving a broader range of collaborators in the innovation creation and leveraging process, the quadruple helix has the potential to offer a multitude of outcomes, which are not limited to solution-oriented innovation. Hence, not only **without stakeholders’ participation, the LL experience would fail** but a degree of reflection and planning should be taken for their selection and gathering since the number, background and the stakeholders’ initial interest will be one of the governing factors that impact the knowledge share, innovation leveraging and collaboration obtained from the KIE program.

### 4.4. Framework strategies for end-user engagement and value creation: The context determines the approach

LLs have emerged, as spaces for innovative and participative research, for the development and activity deployment by using multi-disciplinary methods and approaches and bringing people together in social contexts around a range of themes [92]. Although the task of engendering end-users interest to collaborate with the LL program can, indeed, be considered uncomplicated (as previously stated in section 4.3), it is also universally recognized [1, 4, 75] that the more intricate challenge lies in actively involving them, in a tangible manner and subsequently fostering their sustained participation. While Ahmadi et al. [75] reported challenges associated with schedule compromise between both organizations and individuals, Compagnucci *et al.* [1] highlighted the difficulty in involving the stakeholders in a practical way in the LL activities. In this research study, the TheBrainLL facilitator reported the adversity felt in getting all the parties (namely end-users, providers and host) in the same session. The schedule incompatibilities were sharpened when health professionals’ end-users were considered. As stated by a facilitator “More than the interest in participating, in my opinion, is the time specialists have to dedicate to this type of activities”. To overcome this challenge and guarantee the meeting of interested providers and end-users, a higher number of engaging sessions were developed. Also in TheBrainLL a strategy of developing the ‘Engagement with end-users session: healthcare associations and translational medicine company’ session on a Saturday morning to avoid jeopardizing end-users weekly working schedule and the development of the ‘Engagement with end-users session: Neurology healthcare professionals’ session during the Hospital neurology department monthly general meeting, was adopted. Additionally, to compensate for the engagement sessions’ limited time, after sessions interaction between both parties (providers and end-users) was encouraged. RemotMonitLab shared the struggle associated with the limited time availability demonstrated by end-users to participate in the sessions. As stated by the facilitator “it was difficult to find end-users available to participate”. This challenge has also been reported in literature [58] as LLs deal with the complexity of having multiple stakeholders, with different (and sometimes even opposed) agendas and interests.

It is pivotal to develop LL activities/sessions that motivate end-users to actively participate in the ideation and/or innovation process while also showing and allowing end-users (and other stakeholders) to capture value from the collaboration with the LL program (partners’ value creation will be discussed in detail in the next section). As a strategy to empower end-users Ignasi Capdevila [58] suggest the offering of support and guidance while confronting the resistance towards participation and engagement during LL activities. In a complementary strategy to end-user empowering, we suggest that it is also essential to consider the end user’s underlying motivation to actively participate in the ideation and/or innovation process as well as their incentive to be part of the LL community. By synchronizing with their motivational factors, we believe it is possible to create a safe and cohesive environment that not only allows for constructive knowledge sharing and consciousness-raising but also the cultivation of a sense of purpose and ownership among end-users that will allow their engagement to be personally enriching. This strategy will also encourage the partnership participation of the end-users within the LL community. As recommended by Anna Ståhlbröst [3], the users involved in the LL activities should be addressed with respect since they invest time to participate in the program, often through a pro bono service.

The symbiotic interdependence between end-users value creation by the LL and the KIE dependency on the end-users involvement and motivation led to the end-users engagement as the cornerstones of the living lab approach. In an attempt to address this challenge, a three-stage framework methodology was adopted for the six LLs considered in this study (see Figure 5). This aims to guarantee the solutions progress in their path to the market while also increasing their level of innovation and ability to fit the market needs which will result from the feedback and active participation obtained from the end-users’ engagement and partnership established with the providers. NDAs were signed when confidential information was shared to increase the level of confidence among the participants.

The first stage, the conceptualization phase, was characterized by a set of ‘familiarization activities’ which were tailored to each LL context. These activities aimed to further the analysis of the participant innovation with a reflection on their practical context and market trajectories as well as the provider’s goals and expectations towards the LL program. Some activities were also aimed at understanding and acknowledging the end-users’ needs and expectations towards the KIE program while also developing a reflection of their involvement, contribution and value attained from their participation in the LL. During the conceptualization stage, the end users did not actively interact with the providers, therefore, no feedback on the innovations from the end users was received. In the PCCLL’s first activity during an online session, the participant innovations were presented by the providers’ team to the oncological patients’ family end-users. This session allowed the end-users to gain an acquaintance with the participating solutions and their teams. The PCCLL’s second action consisted of LEGO® Serious Play® (LSP) activity session. TheBrainLL and RemotMonitLab first conceptualization phase started with a kick-off meeting followed by an LSP activity session. During the kick-off meeting, the facilitator from each LL provided the teams’ with an overview of the program’s contextual background and goals. Additionally, the collaborative host institute/company extended a cordial welcome to the participants. In the SLP apparently paradoxical approach (created by Johan Roos and Bart Victor [93] in the mid-1990s) participants guided by a series of open questions communicate and solve problems using LEGO blocks as mediating artefacts to build symbolic or metaphorical representations of abstract concepts [94]. This way, by creating Lego models participants learn and develop a critical reflection on a subject, as well as, a shared understanding in groups [95, 96]. The three LL program SLP sessions were developed with a total of 5 to 6 providers’ team participants per action (*i.e.* which translated in some cases into the development of two sessions per LL) and were both guided by the facilitator Filipe Santos-Silva [97–99]. This was an interesting approach which allowed for an organized rich discussing between participants and conclusive reflections on the following focus topics: (1) how is the team working interaction and its hierarchical organization; (2) does the team members remain sufficiently aligned with each other, with a common vision and common interests; (3) how can the innovation key feature be efficiently and clearly translated to a general public and (4) what are the provider’s expectations towards their participation in the LL experience. The PCCLL, TheBrainLL and RemotMonitLab facilitators all reported the development of comradeship between the teams due to the SLP activity. Both Microbiomlab and AnimalHealthLL opted to develop a kick-off first-session meeting headed by the host. During this session, there was: a welcoming introduction of the purpose, goals and expected outcomes of the LL program, a presentation of the hosting installations, a shared understanding among participants on the lines of communication to be established during the program, an alignment of expectations and objectives for the experimentation plan as well as the definition of timelines and milestones, an assurance of the host availability to foster meetings with end-users and also the commitment of the host staff roles in the activities to be developed.

The conceptualization phase also involved a diagnostic and appraisal session between the Resolve-Health facilitator team and each provider. This activity was present in all the LLs considered in this study. This session involved discussing with each provider the cost to be covered in the project, budget allocation and other complementary possible funding sources. Hence, during this session, the projects’ specific requirements for the innovation leveraging into the next-generation solution level, the assessment of the provider’s degree of commitment and how the LL program can effectively fulfil those requirements, were discussed with the reaching of an agreement.

The second stage, the concretion phase, initiates with the definition of a starting point action plan constructed from the information gathered during the conceptualization phase. Despite this initial action plan role to provide a guideline for the first end-user engagement activity/session, the following concretion phase activities were devised based on an action research (AR) methodology. The PCCLL, TheBrainLL and RemotMonitLab first focus groups were characterized by the resource to the SWOT analysis tool for the discussions and brainstorming with the end-users on the provider’s innovation strengths, weaknesses, opportunities and threats (Figure 5 sessions with a filled circle). Despite the claimed subjectivity associated with the traditional SWOT approach qualitative analysis when applied to strategic business planning [100] and its incapacity to translate the complexity of organizational or business structures [101], when applied to LLs end-user engagement sessions this methodology has demonstrated to be a simple valuable tool. By contrast, to the common adoption of this methodology to analyze LLs ecosystem [102–104], to the authors’ best knowledge, the PCCLL, TheBrainLL and RemotMonitLab LLs’ were pioneers in the application of the SWOT framework as an end-user engagement tool. An openness and acquaintanceship were verified between the participant teams during TheBrainLL’s first end-user engagement session. The bonding established between the providers led to a free exchange of ideas and mutual aid between them. It was also interesting the verify that this activity even led to a collaborative *in vivo* experimentation between two teams. The ThebrainLL facilitator described SWOT as being a ‘useful engagement tool that stimulated opinion sharing and debate’ and also highlighted the advantage of the underlying SWOT concept being perceptible for both the providers and end-users from different backgrounds. A remark was also made by the ThebrainLL facilitator on the future improvement of the sticky notes and the system of pasting them into the corresponding categories. She suggested the replacement of this regime for a digital filling of the SWOT categories by the end-users (*e.g.* in their laptops). The suggested alternative is environmentally sustainable, less time-consuming and simplifies the facilitators’ feedback gathering from the end-users. In the RemotMonitLL, a more intimate setting was developed by the reduction of the number of total participants involved in the SWOT actions to improve the quality of the interaction between the providers and the end-users. This strategy also increased the time available for interaction between the end-users and the providers. The RemotMonitLab facilitator described the SWOT methodology as being an icebreaker that helped the participant share their perspective despite the difficulty felt by some providers in communicating their solutions to the end-users. This facilitator also shared that during the ‘Alignment with end-users and specific stakeholder session’ both providers and stakeholders actively participated in the action which led to the establishment of vital collaborations and the interest of some end-users in maintaining a long-term interaction with the providers. In general, the facilitators from the three LLs reported that they felt well prepared during their concretion phase first session which was characterized by a reduced occurrence of manageable unexpected situations. There was also a general agreement between facilitators that the provision of a coffee break (*i.e.* light snacks and beverages) created a “space” for informal conversations and the development and reinforcement of bonds between the participants. The benefit of having a coffee break to increase the interaction between the participants was also confirmed by Irma Mäkäräinen-Suni *et al.* [12]. We would also like to address the contentious topic of having online sessions. According to the literature [1], real or a virtual environment did not affect the ability to reach the stakeholders. There was, however, a common criticism made by the PCCLL, TheBrainLL and RemotMonitLab LL facilitators which underline the isolation experienced by the lack of face-to-face interaction of end-users that attended the sessions online (*e.g.* due to geographic location). This contributed to their reduced participation and difficulty in keeping them focused and motivated. As stated by the PCCLL facilitator the online sessions ‘jeopardize the flowing exchange of ideas and alternatives were adopted to replace them’. Opposite statements were registered among the providers (Table 5) which claimed “It was useful to have the sessions in an online format” and “I would suggest that all sessions can be assisted via Zoom. Some people live on the Islands, which makes it difficult to travel”. Such disparities in opinions reflect the different perceptions on this subject. Thus the providers give the utility point of view regarding the availability of having online sessions whereas the facilitators provide a perspective on the loss of engagement as a result of the online sessions. Hence, it is possible to understand that although the virtual sessions are usefully tools for connecting partners and developing actions which otherwise would be impossible, it is important to handle this resource with caution and the advantage/disadvantage weighting factor should be taken into consideration. Overall, the facilitators verified an interesting degree of involvement between the providers and end-users which can be associated, not only with the development of tailored sessions based on information collected from the conceptualization phase but also due to the adoption of the SWOT engagement tool.

As previously indicated the following actions in PCCLL and TheBrainLL were designed based on AR. This research approach (first introduced in 1946 by Lewin [30]) aims at both taking action and creating knowledge or theory about that action as it unfolds [105]. Hence this methodology strives to understand complex human environments and processes, obtain information about particular situations, introduce changes, observe the effects of their intervention and enhance scientific knowledge by developing new models and theories [75] and producing guidelines for best practices [106]. Although AR emerged as an established social sciences research method, its effectiveness was also demonstrated in the LL context, as shown in previous research [107, 108]. According to Schaffers *et al.* [109], the strength of the alliance between AR and LLs is due to the KIE’s openness and cooperation, complex social processes, and the need to introduce changes into these processes and observe the effects during the process. To stimulate end-users’ openness, tackle the changing engagement between the participants and increase their value creation, the sequential sessions developed in PCCLL were improved or designed with the feedback obtained from previous session participants’ feedback and the facilitators’ analysis, in a cyclic approach. The actions developed were:

- ‘Survival war stories’ where end-users shared their survivor testimonies with an underlying solution-based focus attitude. During this session, the end-users participated in the problem-solving operation by playing an active role in the improvement and optimization of the innovations;
- ‘Brainstorm sessions’ were meetings where specific end-users and experts discussed innovations’ leveraging particularities;
- ‘Outcome session’ where the developed prototypes and evolution accomplished by the innovations were presented to the end-users.

The PCCLL facilitator emphasized not only the contribution of the heterogeneous group of end-users with different life stories, perspectives and backgrounds for the success of this LL but also the significance of the tailored AR-based session which prompted immersive discussions and creative feedback on the technologies and allowed the providers to collect substantial and impactful opinions. The facilitator also underscored the benefit of conducting multiple rounds of feedback not only in the quality of the engagement but also in the development of a trustworthy and hospitable environment. It is important however to highlight that despite the excellent end-user engagement obtained in the PCCLL, some end-users refused to share their views. As stated by the facilitators ‘despite the activities conducted were considered interesting, they preferred to use that time to listen to others and be in the community’. This illustrates that despite the desire and effort to involve end-users, their engagement in some cases can remain limited. The facilitators from PCCLL also reflected that for the improvement of similar LLs, it would be interesting to include in the concretion phase a wider network of end-user intervenient namely oncological patients themselves or cancer survivors, other caregivers such as oncological children families’ which many times provide their primary source of physical and psychosocial support [110] and also teachers, schoolmates, and peers which are responsible for patient’s psychosocial adjustment intervention [111]. In TheBrainLL and DigitalHealthLL’s ‘Engagement with End-users session: Neurology health professionals’ focus group was re-designed based on the previous session outcomes and data obtained (facilitator records and participants feedback). Hence, in this session, the SWOT analysis was replaced by the Start-Stop-Continue methodological framework as an end-user engagement tool for structuring conversations and stimulating opinion sharing. This methodology is regarded as an interesting simple tool that elicits honest input in team discussions which personnel from diverse knowledge levels and backgrounds [112]. The Start-Stop-Continue framework allowed the evaluation of the innovations by soliciting the health professional perspectives on what they understood the providers should: continue to improve or retain which will spur the reflection on the novelty strengths (continue), which allow for the introduction of new ideas (start) and no longer do which will enable the warning of the solutions’ weaknesses (stop). To guarantee the discussion flow, the facilitators were responsible for the collection of the feedback provided. According to the facilitator, ‘there was no need for my intervention to feed a debate which prospered easily and engendered a rapt enthusiasm and participation of the end-users’. Also as a token of appreciation from the facilitators for the end-user’s availability in developing this session during the Hospital Neurology Department’s monthly general meeting and to enhance the establishment of relations and promote collaboration, an after-session lunch was prepared for all the participants. The midday meal event allowed for the exchange of contact between participants and also the commitment from some end-users to support the validation of a solution in a hospital environment. Hence this session allowed not only for meaningful engagement but also for the achievement of effective feedback, the emergence of a collaboration between participants and the demonstration of interest by the end-users in the realization of a similar session. TheBrainLL also had ‘One2one meetings with specific experts’, more precisely specialized consultants with profound knowledge in patent evaluation and others in patent writing. The latter helped the providers understand the viability of a patent submission on their innovations and how this decision could in the long term impact the innovation growth strategy and in creating value. All the facilitators reported the AR to be a key strategy in providing feedback for the constant adjustment of the sessions to the provider’s needs and the end-user’s particularities and motivations. They also claimed that the AR tool was paramount, during their role as action developers and designers, by allowing the constitution of an interactive learning process which allowed for a constant improvement of the sessions’ quality. In a more inclusive approach to AR, named participatory action research (many times considered as interchangeable terms) is suggested by Logghe and Schuurman [4]. In this study, the involvement of end-users at the macro level design and organization of the LL end-user activities and operations was evaluated. It was found that end-users were able to quickly gather issues and frustrations on one hand and rapidly co-create and implement practical solutions on the other. This is an interesting approach that leds to the arising of the question on how more suitable and engaging the concretion phase sessions of these LL’s would have been if the end-users actively participated in the designs and organization of the KIEs activities.

In the concretion phase of PCCLL, TheBrainLL and RemotMonitLab involve a sequential and complementary series of focus groups with the end-users that not only allowed for the solutions’ evaluation but also the establishment of collaborations and preparation for the testing/validation of the innovations. In contrast, the remaining KIEs concretion phase is characterized by a continuous solution validation program in DigitalHealthLL or residency program in the case of MicroBiomeLab and AnimalHealthLL that allows for the direct testing/validation of the innovations in a real setting and periodic or isolated contact with end-user during that process. Notably, in DigitalhealthLL, MicroBiomeLab and AnimalHealthLL the host institution, the hosts have a central role in the activities’ operational design and coordination. Namely by taking in the providers in their installations, allowing the testing/validation of the prototypes on their internal end-users (*e.g.* patients) or in their settings while also managing and designing the meetings with internal end-users (*e.g.* physicians) and experts (*e.g.* computer engineering technicians). In the case of DigitalHealthLL, the host (CUFF) is a private hospital and it is allowing the testing and validation of the digital solutions prototypes on their patients and with the support and supervision of their health professional staff. Hence, the providers will have feedback as well as the participation of the end-users in the improvement of the technologies. Biocant, the MicroBiomeLab host, is in the concretion phase granting their installations for the testing and/or validation of the provider’s innovations in a real-life setting. They are also providing access to their close end-users network and a set of experts such as Genoinseq [113] (a multidisciplinary team of specialists that offer expertise/services in metabarcoding, functional metagenomics, Human microbiotic, DNA extraction and library preparation and sequencing on Illumina platforms) and other partners within the Biocant ecosystem. In the AnimalHealthLL the providers are testing and validating their innovations in animal patients from the OneVet Porto Veterinary Hospital facility and under the supervision of the Clinical Director and the animal health professional’s staff. This LL is integrated into a network of proximity veterinary clinics and nationally referenced veterinary hospitals which have an internal sharing of cases system at the national level. This broadens the number of animal patients that can be included in the test/validated study and allows the development of more complex evaluations. Due to the extensive required duration of some studies which could go up to 4 years, the DigitalHealthLL, MicroBiomeLab and AnimalHealthLL concretion phase validation program/residency action is still in its development phase. However, we can ensure that the activities are ongoing as expected as the solutions have the management and end-users support system and resources required available.

The last stage, the specialization phase, consisted of a series of interlocked evidence-based and practical guidance training sessions delivered by experts from IP protection, Health/Medical Software regulatory affairs, legal aspects in data protection in health technologies legal aspects in the incorporation of companies, marketing and branding, supporting tech SME and start-ups, national innovation incentives and venture capital knowledge sector. Different learning and engagement frameworks were considered to guarantee an interactive approach between the providers and experts. The sessions’ dynamics were tailored to the number of attending participants, the subject being addressed and the expert level of interest and motivation. The providers from all six LLs’ were invited to participate in these shared training sessions which were not mandatory. In the ‘IP protection’ and ‘Bridging with VCs’ sessions the experts provided a lecture in their field of expertise which was followed by one-to-one training with the providers where project-specific strategic advice and clarifications were given. The other sessions followed a type of course framework with different formats (on-site, hybrid or fully online) where there was a detailed lecture on a certain subject or a set of themes to pass knowledge to the providers. A participatory interaction was encouraged and the providers were spurred to collectively share their questions and their project case for counselling and discussion. The shared sessions between providers were a viable strategy to guarantee their access to invaluable key knowledge for innovation progress and the introduction of the solution in the market. As stated by two participant providers (Table 5): “This is a highly relevant leverage program. I highlight four aspects: … 3) complementary training via workshops/seminars and contact with stakeholders” and “The speakers were relevant and knowledgeable, and the sessions were interesting.” The shared training sessions between providers also valued the experts’ time by sparing them a lecture repetition. The training sessions allowed for the gathering of the best suitable specialists and connoisseurs who were not only able to provide key insightful information that could catapult several innovations development and their introduction in the market but were also able to provide personalized case-specific advice when requested. However, it is important to highlight again the bashfulness felt by the providers that attended the training sessions remotely and which absence of personal contact constrained them to ask questions during the courses. It would have been interesting to arrange a private chat session with the experts to tackle the isolation felt by the online participants. The specialization phase also comprised a final event, the WARM (abbreviation for Worldwide Accelerators Rally at Matosinhos) that aimed to foster a collaborative movement of driving innovation in healthcare. This event brought together innovation acceleration programs, incubators, Technology Transfer Offices (TTOs), investors, companies, and startups from the regional and national innovation ecosystem. This enlightening and inspiring event allowed for the LL’s participant solutions to be presented while allowing the providers to have access to valuable insights, inspiring talks, and to develop collaborations and expand their network of stakeholders.

In this study, it was possible to verify that there are no standardized actions that can be generally adopted for all LLs in the health domain since **the context determines the approach**. Meaning that the complexity of the context in which the innovation will be implemented due to the diversity of projects involved, their needs, and the participant’s contributions and motivations require the development of customized activities. Notwithstanding, an organizational framework based on three phases (conceptualization, concretion and specialization) to improve the KIE’s actions, build trust and increased value creation for all parties and spur end-user engagement, was proposed. Within the three-phase framework strategies, several engagement models and tools (*e.g.* SWOT) that could be adopted and improved by similar LLs, were also shared and discussed. We also highlighted the AR potential as a tool to iteratively experiment with new ways of involvement by collecting data, reflecting on it and planning for upcoming activities. Hence, this method allows simultaneously tackling the complex LL social ecosystem by improving the quality of the LL actions as well as learning about the specific social system and the general expansion of scientific knowledge regarding KIEs. We, however, do believe that the issue of how to optimize LL’s actions could still benefit from further research. In this study, the facilitators developed the role of action researchers. Here they continuously assessed and correspondingly designed/adapted methodologies to ensure the innovation’s progress based on the understanding of the solutions, providers’ and end-users’ needs and variable feedback obtained from all the collaborating participants. As a complementary improvement to AR, and as future research, we suggest the evaluation of the benefits of having all the parties involved in the LL program (host, providers, and end-users) directly participate in the design of the LL actions with the facilitator. Having a representative from each party in the improvement of the LL actions could offer a unique perspective based on different participant expectations and experiences, enabling the adjustments to address their unmet and unaddressed needs by involving them in the problem diagnosis, action intervention, and reflective learning. This strategy could also provide additional empowering and value creation to the KIE participants due to their more active participation and new important role.

### 4.5. Value creation: The most valuable deliverable of LLs

The LLs’ processes support value creation in two general distinct ways, for their KIES partners (*i.e.* providers, hosts and stakeholders) and the presumptive customer of the developed innovation [3]. The concrete conditions under which LL partners get to collaborate and their role in the engagement activities will imprint the process of value creation. Therefore, it is important to understand the roles played by the LL partners (but mainly by end-users) as innovation drivers and consequently the value they provide. In PCCLL, TheBrainLL and RemotMonitLab (Figure 6), the users signalled their needs, shared experiences, evaluated the impact of the solutions, shaped the development of the innovations, supported the innovations validation and expanded the provider’s collaborator network. These contributions led the end-users actions to fall into two considered active roles as an advisor (feedback provider to improve the innovation or any of its specific characteristics) and/or a tester (supporter or participant in the testing of the innovations in a different set-up) and/or a passive role as connector (liaison for the establishment of a connection to a new collaboration with other stakeholders, *e.g.* by introducing or providing the contact of an external stakeholder that can contribute to the innovation progress). As demonstrated in Figure 6, the advisor was the most prevailing role in the TheBrainLL and RemotMonitLab, contrarily to PCCLL where the tester role was predominant. However, it is important to highligh, as stated by Lipp *et al.* [87], this participation is not binary being either active or passive, it comes in different types, some more active and others less so. In the LLs studied, the same end-user could assume multiple roles. As an example in the RemotMonitLab, the psychiatrist end-user not only provided professional advice but also allowed himself to be a test subject of a solution. There was no co-creation (denoted as the process by which these actors come together and the mode of collaboration between them [87]) in any of the LLs considered in this study. Hence, the end-users did not assume the role of initiators (collaboration in the ideation process) and developers (development/building of innovation). As prior research suggests [114], the evaluation and testing are more a less taken for granted as the end-user role while the active end-user participation through co-creation is regarded as an ideal practice in the LL research. Experts assumed an advisor and/or scouter role (looking for innovative solutions or clients). The same roles were adopted by the hosts. All the KIEs participant’s degree of involvement in the LL’s actions/activities and the support and contribution they provided for the innovation process resulted in outcomes obtained during the program which consequently reflected directly in value creation for some of the LL partners. The considered outcome metrics were: ideation (*i.e.* the creative process of generating, developing and communicating new ideas for a new product/service development [115]); innovation evaluation (*i.e.* improvement of the innovation based on the feedback obtained by the stakeholders), innovation development (*i.e.* progress in any specific innovation feature for example software development), validations conducted (*i.e.* the supporting or corroborating on a sound or authoritative basis of its worthiness or legitimacy [116]), testing undertaken (*i.e.* assessment of the innovation performance to new unseen data), prototyping (*i.e.* prototype development) and commercialization strategies elaborated (*i.e.* strategy to help the innovation reach the market for example through the development of a business plan). As it is possible to verify in Figure 7, solutions validation was achieved in all the LLs considered in this study and corresponded to 40.4% of all the outcomes achieved in the ResolveHealth LL program. Hence value was created for the providers (and the experts/host LL which have an innovation scouting role) through the effective and rapid validation and when applicable development (10.6%), prototype creation (8.5%) and innovation testing (2.1%). This value was only acquired with the participation in the LL program and that translated into the innovations’ additional progress to turning into a tangible and impactful product to be introduced in the market. There were no registered ideation and co-creation outcomes since the end-user did not take in the roles which would lead to those initiatives also these outcomes did not fit the innovation requirements goals. There was also no commercialization outcome since the LL’s participant innovations still required significant progress and technical leveraging which hindered at this stage their market introduction.

There was a transcendental benefit for all the participants which consisted in the value creation from knowledge exchange. The knowledge fostered between parties is derived from shared experiences and know-how and discussion/debating on different points of view. The knowledge acquisition with participation in the LL and its role as a value-generating tool for the participants has gained considerable recognition and its importance validated. As an example, Marina van Geenhuizen [117] has developed a conceptual framework for evaluating an LL where learning was looked at as the main outcome of the process. Although this value created is more difficult to quantify, in this study the provider’s knowledge acquired was considered tantamount to the number of evaluation outcomes obtained. As demonstrated in Figure 7, all the participating teams had access to knowledge since the number of evaluation outcomes is considerably proportional to the number of participating projects in the LLs. Although evaluation outcomes were only registered in PCCLL, ThebrainLL, RemotMonitLab and DigitalHealthLL it is almost certain that they will also be present in MicrobiomeLab and AnimalHealthLL. However, at this stage, there is no information regarding the precise number of solutions that will have evaluation feedback from end-users.

Even though they could not be quantifiable, other values created that could be extended to all the LL’s partners are power and empowerment [75], bonding developed from cross-interactions, consciousness-raising [75] and education [75]. The power gaining and the increase in their sense of empowerment could be obtained from both the learned knowledge (acquisition of innovation tools know-how and learning culture [13]) and the recognition of their authority and impact of their contributions. As stated by Anna Ståhlbröst et al. [3] there is an emerging trend of end-users to influence products and services that they might use. In the Resolve-Health program, empowerment was observed with the MicroBiomeLab, DigitalHealthLL and AnimalHealthLL host coordinators. In these LL cases, the host is a key element in the determination of the LL actions and the end-users involved with the providers. Since they have a critical power in determining the ecosystem in which the innovation will integrated and consequently their outcomes. Another example was reported by the PCCLL, TheBrainLL and RemotMonitLab Another unmeasurable benefit from participation in the LLs is the business value created. This is impactful mainly for the providers and some hosts and end-users/expert partners with the reaching of long-term prosperity and growth which is of vital importance for their survival [3]. In ResolveHealth, the providers experienced this value by being involved in the LL process which provided the resources that allowed the progress of the innovation to evolve in its path to reach the market. An interesting example was the TheBrainLL where some providers’ participation resulted in licensing patents or launching start-ups. For the case of the host institution, we would like to share the business value acquired by CUFF (*i.e.* DigitalHealthLL) and OneVet (*i.e.* AnimalHealthLL) in which participation in the LLs allowed them not only to support new entrepreneurial initiatives but also to catalyze and possibly adopt the next great solutions. Although some experts (such as the VC, for example, with their participation in the “Bridging with Investor” session) created similar values to the host institutions, other experts had value creation by finding future customers for their companies through their participation in the LL program.

To attain an additional quantification of the value created by the providers’ participants in the LL their response to the closed answers to the questionnaire was taken into consideration (Figure 8). Hence, data were considered regarding the relevance of the sessions and mentorship, specification of actual needs, final gains and perception of general satisfaction experienced during their participation in the LL program. A total of 13 responses were collected, which represents 33% of the total participants. Three important reasons account for this low adherence to the questionnaire reply. First, the Resolve-Health team did not make this questionnaire response obligatory. This was worsened by the quest for a reply being made in the summer month where some teams might have just not responded due to vacations (which happened to some participants). Secondly, also given that the LL still has continuity, some teams did not feel comfortable answering at this stage of the program. Finally, although the instructions were explicit in asking for each team member to reply to the questionnaire, in some cases we got the note that the team got together to answer the questionnaire once, as a team. By considering the 13 overall responses, the program was rated positively as is possible to verify from Figure 8. There was one team that rated the program very poorly in all items. The item in which they rated the program as a 6/7 was the item referring to the quality and pertinence of the sessions that were provided in an open-access format for all the participant teams. Even though this particular evaluation was poor, once the other responses rate the program very positively, the results are not significantly affected by this negative evaluation.

The value created for the providers and hosts was evaluated based on qualitative data collection (Table 5). As previously stated, only around one-third of the providers’ team members’ participants responded and from those only a smaller percentage responded to the questionnaire open questions. Hence, the employment of more qualitative methods such as open-ended qualitative interviews [8] could have guaranteed data retrieval from a wider spectrum of the provider population and would also allow for the acquisition of additional information that could contribute to the future improvement of the LL experience. From the KIE participant hosts, we would also like to highlight the response of Ana Monteiro from Associação Acreditar, see Table 6, which acknowledges the partnership established with the Resolve-Health LL program and also her avowal of the knowledge value created with their participation in the PCCLL “they gave us very interesting clues”.

The value created in the LL will not only impact the close ecosystem but also shed new light on societal dynamics by triggering organizational culture changes [118] and growing societal awareness of the end-user impact on innovation outcomes [1]. However, as suggested by Fuglsang and Hansen [13] expansion of the KIE benefits to society requires the linking up of the LL framework to the wider shared agendas and socio-technical landscape chances.

As participants become a source of knowledge and ideas, there should be an appropriate reward and incentive system to ensure that human values and priorities are advanced [14, 57]. Thus, value creation for all the collaboration partners should be **the most valuable deliverable of LLs.** Hereof, value generation in the form of knowledge acquisition but also actors empowerment, interpersonal bonding, mindset changing, education, and business value and when possible value to society should be created and acquired in a network of cooperating partners that comprise the LLs ecosystem.

### 4.6. The long-term viability of the LL project: Commitment, trust and communication are crucial

The ultimate LL goal is to actively involve end-users and other stakeholders for a long-term period [9], meaning to prolong the KIE aspects of continuous learning and development not only in the capacity of a project but also beyond its end for a long period. Yet, limited attention has been paid to the evaluation of the criteria(s) that ensure the sustainability of an LL over the long term [92, 119]. In a study Mulder *et al.* [120] suggested the use of the harmonization cube in multiple domains and across several LLs, to facilitate a common ground for sharing the essentials to keep LLs “living”. In more recent research, Mastelic *et al.* [92], suggested the evaluation of the LL sustainability from a business model development strategies (*e.g.* Business Model Canvas) and by the identification of the cost structure, customer segments and the revenue stream. However, as stated by Louna Hakkarainen and Sampsa Hyysalo [121], the long-term collaboration can be laborious and volatile. All the LLs considered in this study were designed to be sustained beyond the scope of the Resolve-Health project. Although it is difficult, if not impossible, to discuss the outcomes of the methodologies adopted to keep the KIEs considered in this study “alive”, it is possible though, to share the actions adopted by the LLs considered in this study in an attempt to increase the LL sustainability at a long-term. Firstly, we would like to emphasize that the sustainability strategy based on a case-by-case tailored design was adopted for all the LLs. The Associação Acreditar (host) will continue to support the technology validation of three innovations that participated in PCCLL. The sustainable operation of TheBrainLL will rely on ongoing support for the participant projects, which will be achieved through regular monitoring. Furthermore, this LL will also assist in facilitating research funding applications for national and European projects through its network; by aiding in one project translation into a startup business development and also by ensuring the project’s long-term viability of intellectual property protection. RemotMonitLab will be kept active by monitoring and supporting some projects’ multi-year validation as a result of the partnership they developed with the participating end-users. We would like to share the case of Virtualeap prototype validation in two public hospitals and a private phycology clinic and Nevaro validation in a research institute and with health professional end-user patients. The ongoing development of the DigitalHealthLL, MicroBiomLab and AnimalHealthLL in the Resolve-Healthprogram, as well as, their technologies’ ineluctably long-lasting validation process will guarantee the viability of these KIEs in the very long term. In the DigitalHealthLL, after the end-users and experts’ engagement sessions and beyond the scope of this study, the participating technology innovations will be subjected to an extensive testing and validation program in the CUFF (host) real hospital settings alongside with existing technologies to establish relative effectiveness and accelerate their route to clinical and market adoption. The interest of CUFF to adopt the most promising participant technologies will guarantee this LL’s lastingness in years to come as it will require the innovations’ continued performance improvement over time. During an extensive multi-year residency, participant projects from MicroBiomeLab and AnimalHealthLL will undergo the validation of their solutions at Biocant (host) and Onevet Porto Veterinary Hospital (host), respectively. We would also like to underscore that there was a set of key principles for sustainability fostering which were adopted in all the KIEs. The LLs expected sustainability is driven by participants’ (and host, promoters and stakeholders) commitment as a result of their motivation/interest [75, 122], trust developed from strong relationships between the actors [1, 102] and continuous open communication with all the involved parties [75]. As a strategy to reinforce the relationship trust there is a focus on the building of partnerships where for every LL project, the provider’s team members are invited to become part of the LL community. An effective communication line is also going to be maintained between the enabler team/steering committee and each participant project (and the corresponding host and/or partner end-user) to continue to promote a supportive environment that encourages ongoing engagement and innovation promotion. To strength the interpersonal bond with the LL’s participant providers and stimulate the innovation leveraging, twice a year, official meetings will be schedule and which progress updates and feedback are going to be discussed. Active phone and e-mail communication and regular appointments will also be scheduled with the participant that have ongoing activities in the LL to ensure the outcomes optimization through the response to their needs and challenges and to facilitate the necessary adjustments and adaptations. The contact frequency will be determined by the innovation progress. Also as stated by Compagnucci *et al.* [1] maintaining periodic meetings will allow to enrich and expand all participants’ competences.

We do believe that **commitment, trust and communication are crucial** principles to guarantee an LL ecosystem sustainability. It would also be interesting to tackle each KIE singularity and complexity through the development of a tailored longevity strategy. We believe that the prioritization of clear continuous communication with the LL participants will increase their motivation to progress in the challenging innovation path and to nurture the already reliant ecosystem.

### 4.7. LL actions and results promotion and dissemination: Raise awareness and extend the LL impact

The increased access to computer networks and the increased sophistication and decreased cost of mobile devices are just two factors that have led to the widespread adoption of social media sites, many of which serve as a common platform for people to meet their communication and information needs [123]. Hence, social media was considered a key dissemination tool for the promotion of the LL’s actions and results. To access a wider range of people/public (*e.g.* across different age ranges) multiple platforms were considered for the six LL’s communication namely, Facebook, Instagram, LinkedIn and Twitter. However, only data from LinkedIn was presented since this was the key social media for ResolveHealth actions promotion and dissemination. As stated in the literature [124], this is the most important channel to distribute business-related information and the preferred tool for marketing and management executives to find relevant quality content. The results confirmed that this platform was not only able to captivate the attention and interest of the target audience followers from the research and education field but also extended their followers to the population from the business development and healthcare services. It was also interesting to verify that the demographic distribution of the followers was 61.6% from Porto and Lisbon metropolitan areas. This could be a reflection of Portugal’s tendency to be a very centralized country (one of the most centralized in the Organization for Economic Co-operation and Development, OECD [125]) with a trend towards coastalization [126] and a territorially polarized population where 44.5% of the national inhabitants is living in Lisbon and Porto (as given in the 2021 census [127]). The other 16.3% demographic distribution of the LinkedIn followers were from Portuguese cities located in the north and central region of the country. This demonstrated the existence of a regional localized interest in the Resolve program and its activities. Hence demonstrating a need for future development of strategies to increase the promotion of interest in the other regions such as in the south of Portugal which has an interesting growing innovation ecosystem. Hence, the LinkedIn social media results have shown an outreach of the LL’s information mainly to national wide followers which could be inferred from the total of only 11.7% international followers. The international followers had an higher incidence from Brazil and UK. This values can be explained by the participation in the LL’s program of projects with providers or some members of the teams from Brazil and the UK (namely LuckieTech and MultiTarget-ND, respectively) which could have led to an increase in the number of followers in those specific countries.

By evaluating data regarding each LL-specific activities promotion in LinkedIn (Figure 9), it was possible to verify that PCCLL and RemotMonitLab showed a higher average engagement rate while MicroBiomLab and AnimalHealhLL demonstrated a lower average engagement rate. Since there is no direct relation between the number of posts published and the engagement rate obtained (*i.e.* TheBrainLL published a higher amount of posts but did not obtain the highest rate of engagement), it is possible to infer that the engagement rate was a result from Resolvehealth followers and the general public (particularly from Research field, Education and the Hospital and Healthcare services domain) subject preference. It was also possible to verify, in Figure 9 that although the post with information regarding the LLs’ specialization phase actions had the highest engagement rate this value was also associated with an equally high standard deviation value. This means that there were posts which led to a reduced number of likes, clicks, comments and shares while there were others that had a high level of public interaction. This could be explained by the specificity of the subjects addressed in the posts regarding the training sessions which consequently could have drawn the attention of a smaller group of people against the promotion of the WARM event posts which due to the magnitude of the occasion could have capture the curiosity of a wider audience.

The ResolveHealth LinkedIn level of influence is characterized by the average engagement rate over a period between November 2022 and July 2023 (Figure 10). If June 2023 is not considered, it is possible to deduce that the posts led to a considerably constant level of influence characterized by a 6.23% range of variation in the engagement rate percentage during the studied period. Hence, there is no indication of a direct relation between the number of posts and the level of engagement. We believe that the level of engagement would be related to the content of the post rather than the number of publications. This statement is in accordance with the literature [128] suggestion that a high number of posts, by itself, does not mean that the message is deemed important or interesting by other members of the community. The surprising peak average engagement of 21.34% in June 2023 was heavily influenced by a 41.17% engagement obtained in an appreciation and thank post for the contributors and participants in the WARM event where a considerable number of entities and collaborators were tagged. It is also important to highlight that all the posts published on LinkedIn used hashtags (hyperlinks generated by the use of prefixing a string of letters with a hash symbol, #, to help other LinkedIn users find this content, for example, #innovation or #livinglab [129, 130]) with trending keywords to facilitate the dissemination and interaction with the publications. When available the corresponding participant’s/entity’s LinkedIn tag was also presented so that the content becomes a community artefact around which groups can discuss, interact and collaborate [131]. Short videos as a tool to promote LL’s services provided, innovation acceleration outcomes and specific actions for innovations leveraging in social media were not used as often as expected due to time limitations. However, video tool is considered a compelling approach since it is generally not considered expensive and allow the information to reach a wider audience [1]. It would have been also interesting to development of short video interviews to end-users to bestow their distinctive eminence collaboration, to provide them with means to express their experience and to allow the reaching of their perspective to a wider audience. The high rate of social media engagement and quality and coherence of the information provided was guaranteed by the presence of a ResolveHealth program communication manager that was responsible for the communication text writing and some cases verification (*e.g.* draft text for the promotion of specific events were written by the facilitators) as well as the publications content administration.

Besides social media, the resolve-Health organization’s website was also used to share information about the LLs. The non-installation of a website analytical tool at the time of its launching impaired data collection to only the number of posts published. Although the impact obtained from the dissemination of some of the LL’s outcomes on the resolve-Health organization’s website was not quantitatively measurable, we believe they enhance the exposure of the LLs results and program to a wider population.

Other dissemination strategies for the presentation of the LLs’ results were also adopted. We would like to highlight the publishing of the project findings and suggestions for good practices in an international journal; the presentation of the results through a poster and article at an international conference [132], the summarizing of the findings in the project’s final report and issuing of the LLs program success in a press release. Besides the more general impactful promotion and dissemination of the results to a “wider public” it is important to emphasize that all the LL actions had a doubly beneficial effect by allowing not only the leveraging of the participant innovations but also their promotion in a more closed target group. A prime example was the “Bridging with VCs” session which allowed VCs and a Business Angel to be aware of the outcomes reached by the innovations during the LL program and also the WARM event where results of the LL program were presented to different parties in the innovation ecosystem. Besides the communication media considered in this study, other interesting dissemination strategies were reported in the literature [56, 65, 133], namely discussing project activities on the local radio, publishing the successes and specifically targeting information regarding the LL programs in the local or national newspaper, publishing project findings on national journals, sharing results to local community groups (*e.g.* presentation of innovation in market-stands) and through a continue updated newsletter on the LL’s actions.

While there are no standard solutions, it is clear that digital tools are a decisive asset to **raise awareness and extend the LL impact**. This study was able to demonstrate that the principle of maintaining consistent communication through social media can guarantee the promotion and involvement of the general public in the LLs’ actions and their access to the final results obtained.

### 4.8. Environmental sustainability: The goal of having a greener planet

Rapid modernization and industrialization have led to an accelerated deterioration of the natural environment [134]. This presents an urgent need for the development and adoption of actions that ensure the continuity of the earth’s ecosystem’s ability to sustain life [135]. In an attempt to neutralize and reverse the damages done to the environment, there is worldwide progress in imposing standards of environmental accountability [136], in the promotion of green innovations [137] and in the promotion of sustainable entrepreneurship (defined as the discovery, creation, and exploitation of opportunities to create future goods and services that sustain the natural or communal environment and provide development gain for others) [138]. Environmental sustainability awareness is an increasingly relevant aspect of LLs. This has been achieved through the development of LLs that support environmentally sustainable innovations, processes and ecosystems [139, 140], as well as, considering the social and economic impact that the innovation being supported might have once implemented in society [3]. In the Resolve-Health LL program, the environmental sustainability concerns were addressed by implementing environmentally friendly measures. Hence, unnecessary travel (by air or land) was reduced and consequently the associated energy consumption and pollution. Information technology (IT) support tools were also adopted for the development of online LL sessions meetings and communication. The latter was achieved through the online promotion and dissemination of the LL actions and results, the recurrent use of e-mail and phone calls for correspondence, the creation of WhatsApp Messenger online discussion groups (*e.g.* between the Resolve-Health team members), the development of online surveys and the award of digital end-users appreciation certificates.

The stakeholders’ and enabler’s feedback was obtained from informal conversations with the liaisons who were constantly encouraging them to share their opinions, concerns and suggestions regarding the LL program they are collaborating with. This information was registered by each facilitator and considered as a result of the improvement and design of the following LL experience.

In this section, we highlight the good practice of the LLs’ program being concerned with minimizing their environmental impact and adopting as many environmentally responsible practices as possible to contribute to the societal shared **goal of having a greener planet**.

### 4.9. Results information processing: Gathering of realistic and insightful information for the LL performance analysis and future improvement of KIEs

Reduced attention has been paid to KIEs evaluation criteria and how they contribute to the performance assessment [92]. Notwithstanding, there appears to be an agreement on the advantage of employing a diversity of tools for the LL results evaluation [11]. In a recent interesting review study developed by Bronson *et al.* [8] it was claimed that several scholars are recommending different types of models for structuring LL evaluation, but largely for those using quantitative data. He also verified that these quantitative methods are mostly combined with qualitative methods (as a mixed approach to data processing). The data processing methodology adopted in this study (*i.e.* metrics, statistics analysis methods or frameworks) also followed this line of reasoning where both qualitative and quantitative features were selected to effectively address the characterization of LL and the proposal of good practices as key performance indicators. The results obtained were categorized into five groups: providers, stakeholders, engagement strategies, value creation and results dissemination. This strategy helps to organize and assess the information for the evaluation of the LL performance. In ‘providers’ data, information regarding the participant teams and their solutions was assessed through both qualitative (*i.e.* participant projects acronym, short description, TRL and technical support provided) and quantitative (*i.e.* percentage distribution of the participant project in the total LL per innovation filed of application; number total number of innovators and their distribution per gender in each LL) methods. Quantitative data regarding the total and per LL number of stakeholders and the number of end-user by background type as well as qualitative data regarding their background were considered in the ‘stakeholder’ category. In the ‘engagement strategies’ results, only qualitative data was considered due to the AR approach adopted. In this category, each LL action within a three-stage configurational process was presented and described. The ‘value creation’ data gathers all information that will allow to evaluate and infer the quality and amount of value created for all the collaboration parties in the LLs. Therewith, it was considered the end-user roles and correspondent percentage contribution, type and the number of outcomes from the innovation leveraging as a result of the provider’s participation in each LL (both cases mixed-method data processing), the providers’ team close (quantitative data) and open (qualitative data) portion of the survey’s results and also the feedback from the enablers collaboration in the LL’s (qualitative data). The information regarding ‘results dissemination’ was all statistical quantitative data. Although quantitative methods are more common in assessing LLs focused on technology development and technology adoption [8], in this study both qualitative or/and quantitative methodologies were considered and were concurrently (as complementary strategies) or individually chosen to guarantee the optimization in information results processing and to achieve a wide-reaching and deep evaluation of the LLs performance.

The results processing methodology considered in this study allowed for the reliable suggestion of a set of LL good practices, as key performance indicators for the LL’s program’s proper implementation and its ongoing success analysis. To bring further clarity, Figure 12, is an exemplification of the relation between data results processing and their application for the development and evaluation of the key performance LL good practices.

**Figure 12.**
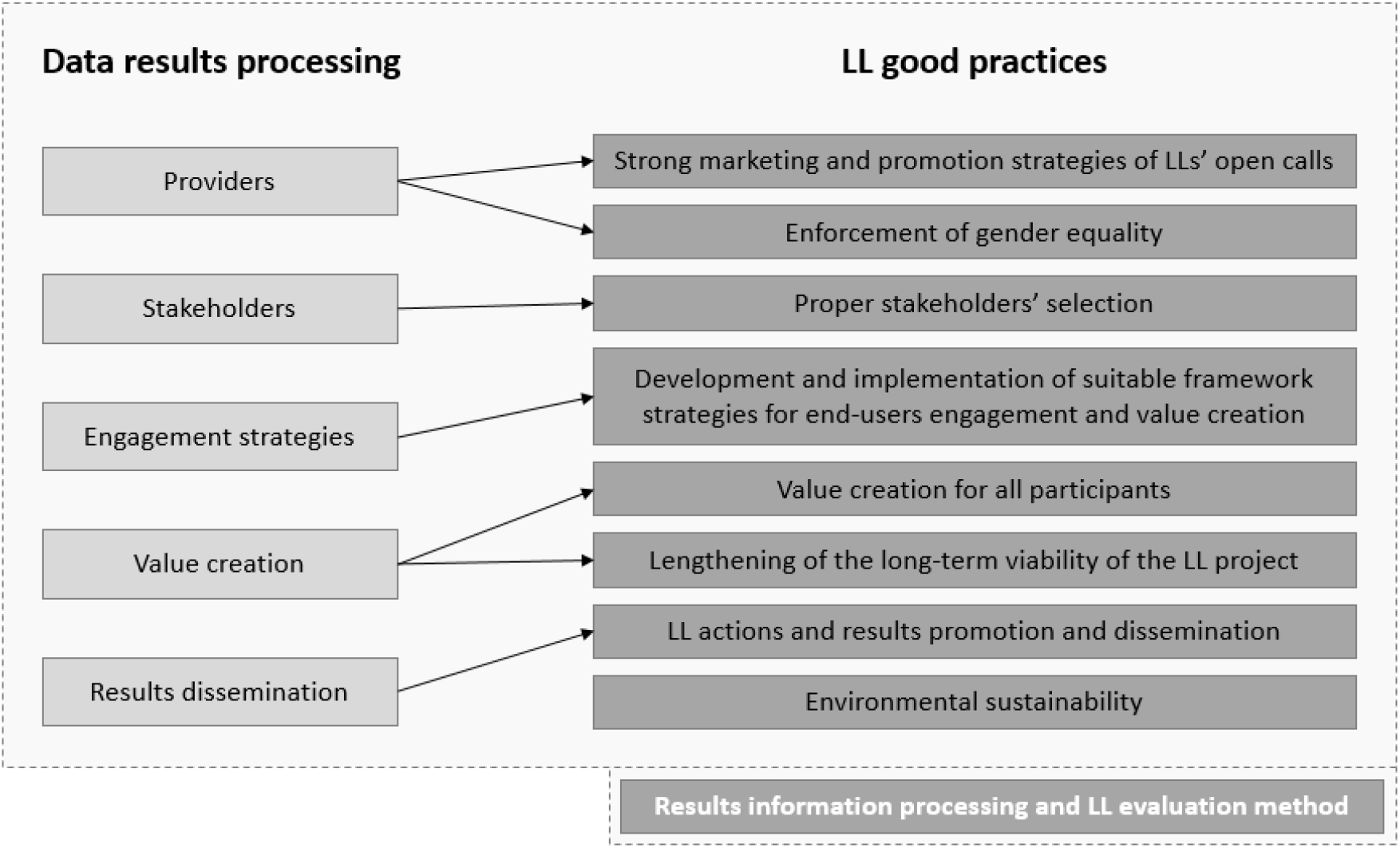
Schematic representation of the relation between data results processing and their application for the development of the LL good practices

The assessment of engagement and diversity of stakeholders/partners/users within the LLs innovation system, were the most common and important indicators of LL success according to Bronson *et al.* [8]. In this study, we propose the use of a comprehensive broad and complementary set of quantitative and qualitative results from **realistic and insightful information for the performance analysis and future improvement of KIEs.** These processed results will not only allow for obtaining more detailed and in-depth knowledge about LLs but also be used as benchmarks to assess the efficient implementation of LL and the development of a set of robust good practices indicators of LL success. The regarded results information, as well as, the data processing and results presentation methods adopted do not have to be limited to the ones suggested in this study since the introduction of additional data and its analysis will enhance the quality of the LL assessment developed. Also, for the LL integral evaluation, it would be pertinent to consider as a key performance indicator, the KIE’s wider impacts beyond the ecosystem created (*e.g.* in society). We recognize the importance of LLs following standardization good practices that can also provide consistency in KIE’s research studies. Hence, we propose a more general but essential set of LLs’ good practices which also present flexibility and comprehensiveness to adapt to the health domain’s LLs complexity and programs diversity. We also like to acknowledge that there is always space for improvement and the standardization feature is time and case-dependent due to these guidelines’ dynamic nature. Meaning that the LL’s good practices should be in a constant process of adjustment and improvement in response to the LL definition and application’s ongoing evolution. In future work, insight into the extended impact of the proposed LL good practices could be obtained with their application to other case scenarios from different geographical areas and which would require the support and leveraging of innovations from different fields of application.

## 5. Conclusion

There are no standardly accepted good practices for health-related LL meaningful and impactful process guidance. Hence, in this article based on the singular and cross-case analysis of six LLs, a more general but essential set of LLs’ good practices for the design enhancement, implementation and assessment of KIEs programs in the health domain are proposed. This contribution is an invitation for the scientific community to consider the proposed set of practices as reference tools to complement and clarify the future implementation and evaluation of their LL studies. The advances made in this article will hopefully also foster further reflection and debate on how to improve the LL’s dynamization and its potential optimization by considering the challenges and opportunities faced by the six LLs considered in this study. From a more general perspective, the data presented will also allow the strengthening and enrichment of the research literature data regarding the constantly evolving and complex LL knowledge innovation ecosystems.

## 6. Author Contributions

Living Lab conceptualization, J.A., A.C., D.O. and F.S., methodology application, J.A., A.C., D.O., F.S., N.R. and S.L.; Writing—original draft preparation, N.R.; image design P.R., statistical study elaboration, analysis and writing, S.L; writing—review and editing, P.H. and M.B.; supervision, P.H. and M.B.; funding acquisition, P.H. and M.B. All authors have read and agreed to the published version of the manuscript

## 8. Acknowledgements

The authors acknowledge the financial support of FEDER – Programa Operacional Competitividade e Internacionalização, in the setting of Contrato de Concessão, under the reference POCI-01-0246-FEDER-181287.

We thank the support from Associação Acreditar as well as the participation and contribution of the “Barnanes”, caregivers and volunteers. We would also like to thank Professor Dr. Marques Teixeira from Neurobios; Dr. Jorge Alves from Centro CEREBRO; Dra. Ana Chagão Monteiro, Dra. Catarina Moreira, Dra. Piedade Sande Lemos and Dra. Maria José Barros from CUFF; Dra. Conceição Egas from Biocant (and Dra. Joana Branco who enabled this collaboration) and Dr. Luís Lobo from OneVet for their participation as hosts in the KIEs. We would like to thank all the stakeholders for their contribution and dedication, without them there would be no LL. We thank Luís Filipe from i3s, for the work developed as facilitator of the LEGO^®^ Serious Play^®^.

