## Supplementary Material Document for "Knowledge Innovation Ecosystem for the Promotion of User-Centre Health Innovations: Living Lab Methodology and Lessons Learned Through the Proposal of Standard Good Practices"

### RESOLVE Application Form

**APPLICANT INFORMATION**

**Promotor (attach biosketch)**

**Name:**

**Affiliation:**

**Telephone:**

**E-mail:**

**Additional members (attach biosketch)**

**Names:**

**Affiliation:**

**Telephone:**

**E-mail:**

**PROJECT INFORMATION**

**Acronym:**

**Title:**

**Stage of development (Y/N):**

( ) Initial data

( ) Full set of data

( ) Prototype made

( ) Validation data

( ) Market and sales data

**Select which area the project addresses (X)**

( ) Mental health

( ) Nutrition for cancer patients

( ) Accessibility to treatments

( ) Patient-clinic interaction

( ) Health literacy

( ) Platforms to ease compliance with school programs

( ) Support for cancer survivors

( ) Digital health solutions

( ) Other:___________________________________

**PROJECT DESCRIPTION**

1. **What health problem and purpose is addressed? (max 1000 characters)**
   *(Describe the problem, stakeholders involved and market dimension addressed)*
2. **Briefly describe your solution (max 2000 characters)***(project/research/invention/idea/technology/product/service/start-up)*
3. **Demonstrate the uniqueness of your solution and how it compares to existing competitors. (max 500 char.)**
4. **Do you have intellectual property protection? If yes, what is the protection status (include relevant dates). (max 500 char)**
   *(Describe obligations to third parties; examples: external sponsorship, investors, contracts, material transfer agreement, grants, software, information or other funding associated to the creation of this technology) (max 500 char)*
5. **Are you interested in launching a start-up to commercially explore the technology? Describe your entrepreneurship skills. ( max 500 char.)**
6. **Are you interested in licencing the technology to third parties? What companies may be interested in your technology? (max 500 char.)**
7. **Explain why you think the RESOLVE-Health 2.0 program could add value to your project. (max 500 char.)**

**Attachments:**

**Cvs (biosketch)** (in one pdf)

**Published material** (manuscript, articles, abstracts, oral presentations) (in one pdf)

**Other relevant information** (drawings, presentations, or other data) (in one pdf)

**Website, LinkedIn** (<add link>)

__________________________________________

**Supplementary Figure 2.** Application form for the LL candidate’s team to apply for the LLs’ program

### Final survey

This evaluation was created so that the RESOLVE-Health Program team could obtain systematic feedback on the quality and pertinence of the activities it promoted with the selected teams. The information collected in “Additional remarks” will be considered to identify the program strengths to adopt in future actions and the weak points to be improved in future editions. The information obtained from this survey will be considered not only for the RESOLVE2.0 final project metrics but also for the development of a research study and consequently results published in a scientific article.

The next items refer to the set of sessions in which your team participated, within the scope of the action for which it applied and was selected. On a scale of 1 to 7, 1 means strongly disagree and 7 means strongly agree.

These data will be analyzed together and treated in a pseudonymized way.

| **Item:** | **1** | **2** | **3** | **4** | **5** | **6** | **7** |
|---|---|---|---|---|---|---|---|
| The program satisfactorily contributed to the development of my project. | | | | | | | |
| The program consistently addressed the needs associated with my project’s development phases. | | | | | | | |
| The mentorship I received was significant for the project's various stages of development that I set out to do in this program. | | | | | | | |
| The sessions that the program contemplated were relevant. | | | | | | | |

| |
|---|
| I managed to achieve the goals I set for myself at the beginning of the program. |
| The information provided by the program helped me to define the needs associated with the development phases of my project. |
| The sessions contemplated by the program helped me identify challenges and solutions for the development and implementation of the product. |
| The financial support provided by the program was relevant to achieving the objectives I set for myself in this program. |
| I managed to exceed the development expectations I had at the beginning of the program |
| The program exceeded my expectations in terms of quality. |
| The sessions that the program contemplated were important. |
| The program coordinators developed sessions with inter-project collaborative environment that benefited the achievement of objectives I set for myself at the beginning of the program. |
| The program exceeded my expectations in terms of relevance. |
| At the end of the program, my level of knowledge was higher than what I had at the beginning. |
| Once the program is over, I leave with a more accurate view of the possible positioning of my product on the market. |

Additional remarks (this free field allows you to add any information that you think is relevant about the set of sessions or any session in particular):

Could you please identify per order the most three relevance the sessions that the program contemplated, according to the characteristics and needs of your project.

**Supplementary Figure 2.** The survey for the participating providers' team.
